# Genes required for the formation of virulence-provoking bacterial sphingolipids

**DOI:** 10.1101/2024.02.16.580706

**Authors:** Roberto Jhonatan Olea-Ozuna, Sebastian Poggio, Ed Bergström, Aurora Osorio, Temidayo Oluyomi Elufisan, Jonathan Padilla-Gómez, Lourdes Martínez-Aguilar, Isabel M. López-Lara, Jane Thomas-Oates, Otto Geiger

**Affiliations:** Centro de Ciencias Genómicas, Universidad Nacional Autónoma de México, Avenida Universidad s/n, Cuernavaca, Morelos, CP 62210, Mexico; Departamento de Biología Molecular y Biotecnología, Instituto de Investigaciones Biomédicas, Universidad Nacional Autónoma de México, Mexico City, Mexico; Centre of Excellence in Mass Spectrometry and Department of Chemistry, University of York, Heslington, York, YO10 5DD, United Kingdom

**Keywords:** bacterial cell envelope, sphingolipid biosynthesis, phospho-sphingolipid, serine palmitoyltransferase, ceramide, dihydroceramide, *Caulobacter crescentus*, *Galleria mellonella*, greater wax moth

## Abstract

Sphingolipids are ubiquitous in membranes of eukaryotes and are associated with important cellular functions. Although sphingolipids occur scarcely in bacteria, for some of them they are essential and, in other bacteria, they contribute to fitness and stability of the outer membrane, such as in the well-studied α-proteobacterium *Caulobacter crescentus.* We previously defined five structural genes for ceramide synthesis in *C. crescentus*. However, other mutants affected in genes of this same genomic region show cofitness with a mutant deficient in serine palmitoyltransferase. Here we show that at least two phospho-sphingolipids are produced in *C. crescentus* and that at least another six gene products are needed for the decoration of ceramide upon phospho-sphingolipid formation. All eleven genes participating in phospho-sphingolipid formation are also required in *C. crescentus* for membrane stability and for displaying sensitivity towards the antibiotic polymyxin B. The genes for the formation of complex phospho-sphingolipids are also required for *C. crescentus* virulence on *Galleria mellonella* insect larvae.

**Author Summary:** Sphingolipids participate in the formation of biological membranes and molecular signaling in higher organisms. Many bacteria also accommodate sphingolipids in their membranes. Here we report that eleven genes participate in the synthesis of complex bacterial phospho-sphingolipids. Our data show that these lipids contribute to membrane stability, but also confer sensitivity towards certain antibiotics. The bacterium *Caulobacter crescentus* is widely distributed in fresh water lakes and streams and was considered to be non-virulent. However, we demonstrate that complex phospho-sphingolipids are the main virulence contributors in this bacterium.

## Introduction

Sphingolipids (SphLs) are essential ingredients of eukaryotic membranes and participate in membrane structural organization, lipid raft formation, cell signaling, and many other important cellular processes (Nelson and Cox, 2021). Although SphLs were considered to be rare components of bacterial membranes, they are more widespread than previously thought (Stankeviciute *et al*., 2022). SphLs seem to be located mainly in the outer layer of the outer membrane (OM) in diderm bacteria and are encountered in many members of the Proteobacteria and the Bacteroidetes (Geiger *et al*., 2010; Geiger *et al*., 2019). The outer layer of the OM is usually covered by lipopolysaccharides (LPSs) which provide an efficient barrier against external toxins and antibiotics (Nikaido, 2003). As LPSs are partially or completely replaced by SphLs in some bacteria, SphLs are expected to also serve as functional replacements for LPSs (Kawasaki *et al*., 1994; Keck *et al*., 2011).

The biosynthesis of SphLs in eukaryotes takes place in five stages (Nelson and Cox, 2021). However, in bacteria only the first step was known, in which serine is condensed with a fatty acyl-thioester to form 3-oxo-sphinganine (3-ketosphinganine) catalyzed by serine palmitoyltransferase (Spt). While Spts of the Bacteroidetes use the thioester fatty acyl-coenzyme A (acyl-CoA) as substrate, like Spts of eukaryotes, recent work clarified that Spts from α-, β-, and γ-Proteobacteria (summarized as *Rhodobacteria*) prefer an acylated specialized acyl carrier protein (AcpR) over acyl-CoA as substrate (Padilla-Gómez *et al*., 2022). The substrate for this reaction, acyl-AcpR, was shown to be formed by a specialized acyl-ACP synthetase AasR (Olea-Ozuna *et al*., 2021; Padilla-Gómez *et al*., 2022). In the α-proteobacterium *Caulobacter crescentus* two more structural genes, necessary for ceramide formation, were identified (Olea-Ozuna *et al*., 2021), one potentially encoding a dehydrogenase (CC_1164) and the other an *N*-acyltransferase (CC_1154). In analogy to eukaryotic ceramide synthesis, we proposed that CC_1164 might be the dehydrogenase that reduces 3-oxo-sphinganine to sphinganine and that CC_1154 might be the *N*-acyltransferase that converts sphinganine to *N*-acyl-sphinganine (dihydroceramide) (Olea-Ozuna *et al*., 2021). A more recent study suggests that the order of these last two steps for dihydroceramide formation might be inverted in *C. crescentus* (Stankeviciute *et al*., 2022).

*C. crescentus* is able to produce at least two different groups of complex SphLs, glyco-SphLs (GSphLs) and phospho-SphLs (PSphLs). Under phosphate-limiting growth conditions, GSphLs are formed by the action of two glycosyl transferases (Stankeviciute *et al*., 2019). For the formation of the PSphL ceramide di-phosphoglycerate three genes were reported to be required (Zik *et al*., 2022).

LPSs as well as bacterial SphLs are synthesized at the inner membrane (IM) of diderm bacteria. However, their final destination seems to be the outer layer of the OM. Transport of LPSs from the IM to the OM has been intensely studied. After assembly of the core lipid A structure in the inner layer of the IM, the MsbA transporter flips the lipid A to the outer layer of the IM where it is further modified (Doerrler, 2006). LPS is then transported through the cell envelope by the LPS transport (Lpt) complex consisting of seven essential Lpt proteins (Bertani and Ruiz, 2018). Initially, the ATP-driven LptB_2_FGC ABC transporter extracts LPS from the IM. Then LPS is positioned onto a transenvelope bridge involving LptF, LptG, LptC, LptA, and LptD. LPSs are pushed across the bridge until they reach the LptDE translocon which accommodates LPS in the outer layer of the OM (Okuda *et al*., 2016). Presently it is not known how SphLs are transported from the IM to the OM.

For some SphL-forming bacteria that lack LPS, SphL synthesis seems to be essential as exemplified in the case of *Sphingomonas koreensis* (Price *et al*., 2018; Olea-Ozuna *et al*., 2021). In bacteria that form both LPSs and SphLs, phenotypes of mutants deficient in SphLs are more subtle than in bacteria devoid of LPSs. In the case of *C. crescentus*, survival at elevated cultivation temperatures is dramatically reduced in mutants that cannot produce SphLs when compared to the wild type (Olea-Ozuna *et al*., 2021). Also, SphL-deficient mutants are much more sensitive to detergent treatment than the wild type (Olea-Ozuna *et al*., 2021). Surprisingly, SphL-deficient mutants are more resistant than the wild type to the antibiotic polymyxin B (Price *et al*., 2018; Stankeviciute *et al*., 2019; Olea-Ozuna *et al*., 2021), maybe due to the fact that they cannot synthesize the SphLs, which might be the target molecule of polymyxin B in *C. crescentus*.

As most bacterial isolates of *Caulobacter* have been isolated from fresh water sources, for a long time this genus was considered to be non-virulent. However, an increase in clinical *Caulobacter* isolates (Justesen *et al*., 2007; Bridger *et al*., 2011) put this perception into question. A recent report (Moore and Gitai, 2020) shows that both clinical and non-clinical isolates of *Caulobacter* show virulence on greater wax moth (*Galleria mellonella*) larvae due to a heat-resistant, cell-associated factor.

Here we show that in addition to the five genes required for ceramide synthesis, another six genes are needed for PSphL synthesis. Mutants in all of these eleven genes are impaired in membrane stability as they are sensitive to the detergent deoxycholate, but at the same time are much more resistant to polymyxin B than the wild type. We also show that intact genes for PSphL synthesis are required for *C. crescentus* virulence on *G. mellonella* larvae.

## Results and Discussion

### Serine palmitoyltransferase required for the biosynthesis of phospho-sphingolipids

Previous studies revealed that Spt is required for the formation of ceramide and GSphLs (Stankeviciute *et al*., 2019; Olea-Ozuna *et al.,* 2021). Applying different thin-layer chromatographic (TLC) separations, we explored whether we were able to detect phosphorus-containing lipids that were formed in a Spt-dependent way. Separation of ^14^C-acetate-labeled lipids revealed that two lipids (Lipid I and Lipid II) were formed in *C. crescentus* wild type, that were not detected in the *spt*-deficient mutant (Fig. 1A). Complementation of the *spt*-deficient mutant with a plasmid containing the intact *spt* gene restored the formation of Lipids I and II, which was not the case when the *spt*-deficient mutant harbored an empty vector instead (Fig. 1A). When the same strains were radiolabeled with ^32^P-phosphate, TLC analyses of lipid samples indicate that Lipids I and II contain phosphorus and are formed in the wild type but not in the *spt*-deficient mutant (Fig. 1B). Therefore, as Lipids I and II require Spt for their formation, they must be PSphLs. Again, the *spt*-deficient mutant with the intact *spt* gene *in trans* formed ^32^P-labeled Lipids I and II, which was not the case when the *spt*-deficient mutant contained the empty vector (Fig. 1B). These data suggest that in *C. crescentus* at least two different PSphLs are formed, Lipid I and Lipid II.

**Fig. 1.**
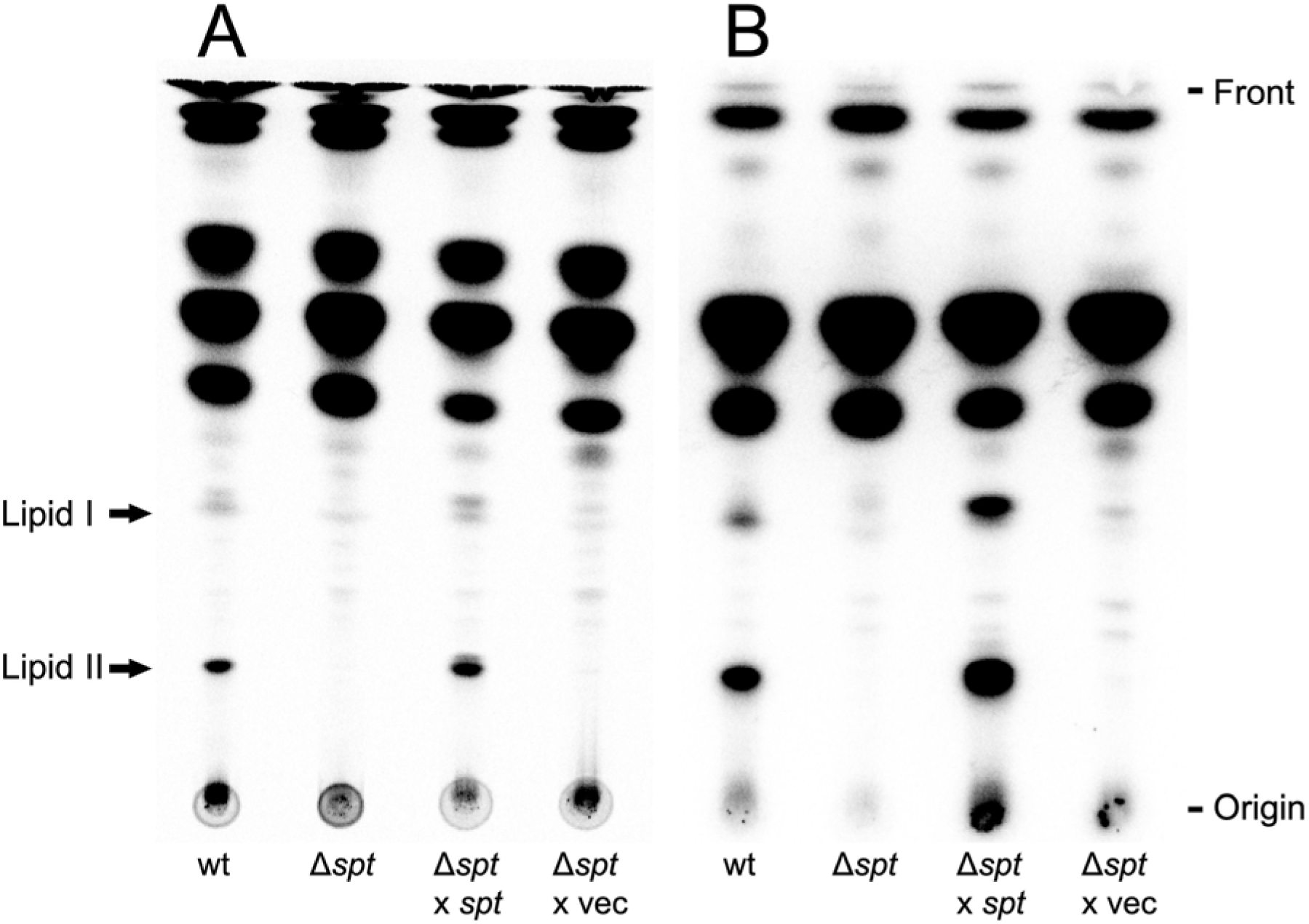
Formation of two phospho-sphingolipids in *C. crescentus* requires the gene for serine palmitoyltransferase (*spt*). Different strains of *C. crescentus* [wild type strain (wt), *spt*-deficient mutant (Δ*spt*), *spt*-deficient mutant harboring the *spt* gene *in trans* (Δ*spt* x *spt*), and *spt*-deficient mutant harboring the empty vector pRXMCS-2 (Δ*spt* x vec)] were cultivated on complex medium in the presence of ^14^C-acetate (A) or ^32^P-phosphate (B). For strains harboring the pRXMCS-2 vector or derivatives of it, xylose and kanamycin were added to the culture media. After harvesting cells, lipids were extracted, lipid samples were separated in chloroform/methanol/acetic acid/water (8:3:2:1) by TLC and developed chromatograms were analyzed by phosphorimaging. Arrows indicate potential PSphLs (Lipid I and Lipid II) formed by *C. crescentus*.

### Ceramide phosphoglycerate formation in C. crescentus is serine palmitoyltransferase-dependent

In a recent study (Zik *et al*., 2022), ceramide phosphate, ceramide phosphoglycerate (CPG), and ceramide di-phosphoglycerate (CPG2) were reported as PSphLs formed in *C. crescentus*. Based on these results, Zik and colleagues proposed a model for PSphL biosynthesis in *C. crescentus* which involved CpgB (CC_1160) as a ceramide kinase, converting ceramide to ceramide phosphate, CpgC (CC_1161) converting ceramide phosphate to ceramide phosphoglycerate, and CpgA (CC_1159) converting ceramide phosphoglycerate to ceramide containing two phosphoglycerate moieties (Zik *et al*., 2022).

On mass spectrometric analysis, our lipid extracts of an *spt*-deficient mutant that expressed *spt in trans* gave a very intense signal (*m/z* 704.48585) in the negative ion mode electrospray ionization Fourier-transform ion cyclotron resonance (ESI-FT-ICR) mass spectrum (Fig. 2A), that was not detected in such spectra of lipid extracts of an *spt*-deficient mutant harboring an empty vector. The signal at *m*/*z* 704.48745 corresponds to an ion of elemental composition C_37_H_71_O_9_NP (theoretical *m/z* = 704.487193, mass accuracy 0.37 ppm), and is thus assigned as M-H^-^ of a compound with elemental composition C_37_H_72_O_9_NP, which would correspond to a CPG. On collision induced dissociation (CID) of the *m*/*z* 704.48745 ion, two product ions were observed (Fig. 2B): an ion at *m*/*z* 166.97503 with the composition C_3_H_4_O_6_P (theoretical *m/z* = 166.975098, 0.4 ppm mass accuracy), which can be assigned as deriving from the phosphoglycerate head group and an ion at *m*/*z* 616.47012 corresponding to an ion of elemental composition of C_34_H_67_O_6_NP (theoretical *m/z* = 616.471149, 1.7 ppm mass accuracy), derived by loss from the precursor of a neutral fragment with the composition C_3_H_4_O_3_ corresponding to loss of the glycerate. We therefore propose that the lipidic compound produced by a *C. crescentus spt*-deficient mutant expressing *spt in trans* is CPG differing from that described by Zik *et al*., in lacking a hydroxyl group on the lipidic portion, presumably the fatty acyl substituent.

**Fig. 2.**
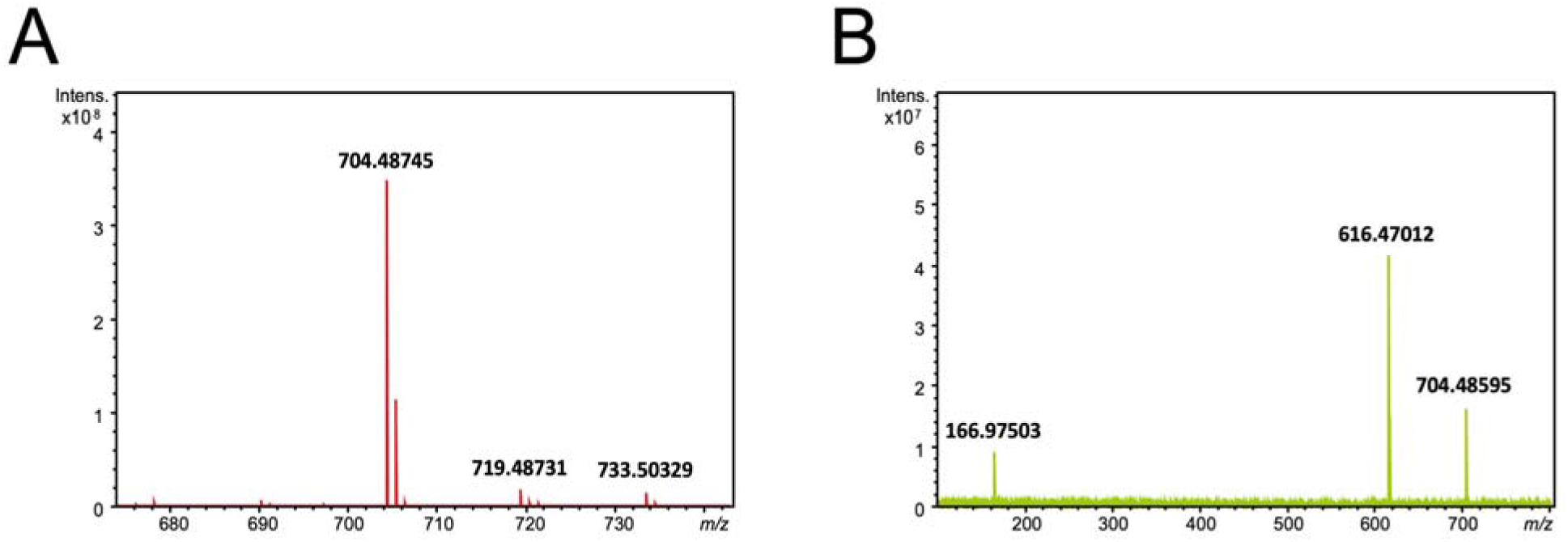
Negative ion mode ESI-FT-ICR mass spectrum of lipid extract of *C. crescentus spt*-deficient mutant expressing *spt in trans*, showing intense M-H^-^ signal at *m/z* 704.48745 (A). Negative ion mode product ion spectrum obtained on collision inducted dissociation of the M-H^-^ precursor ion at *m*/*z* 704.48745 (B).

In order to enrich for Lipid I or Lipid II, we separated by TLC whole lipid extracts from the *spt*-deficient mutant that expressed *spt in trans*, visualized the separated lipids and extracted Lipid I or Lipid II from the silica gel scraped from the plates in the areas in which the compounds corresponding to Lipids I and II migrated, before analyzing them by mass spectrometry. TLC analyses of enriched Lipid I and Lipid II fractions suggested that they had been separated from each other and migrated as two or one defined compounds, respectively (Fig. S1). As a control, neither of these compounds could be enriched or detected after we followed the same approach to purify Lipids I or II from an *spt*-deficient mutant of *C. crescentus* that harbored only an empty vector and therefore was unable to produce SphLs (Fig. S1).

Negative mode ESI mass spectra of extracts enriched in Lipid I and Lipid II (Fig. 3, panels A and D) bear signals consistent with the presence of two types of lipid. Intense signals at nominal *m/z* 704 and 720 are consistent with the *spt*-dependent CPG described above, and its hydroxylated counterpart respectively. The identities of these species were determined on interpretation of their high mass resolution product ion spectra (Fig. 3, panels B and C respectively). In addition, less intense signals were observed in the mass spectra of both the extracts enriched in Lipid I and Lipid II, that correspond to di-phosphoglycerate derivatives of the two ceramide phosphoglycerates. Signals corresponding to these di-phosphoglycerates were observed at *m/z* 872.4709 (Lipid I-enriched fraction) and *m/z* 872.4710 (Lipid II-enriched fraction) for the unhydroxylated species, and at *m/z* 888.4659 (Lipid I-enriched fraction) and 888.4660 (Lipid II-enriched fraction) for their hydroxylated counterparts. The high mass resolution product ion spectra of these two components in the Lipid II-enriched fraction (Fig. 3, panels E and F) are clearly consistent with the presence of di-phosphoglycerate counterparts of the phosphoglycerate ceramides.

**Fig. 3.**
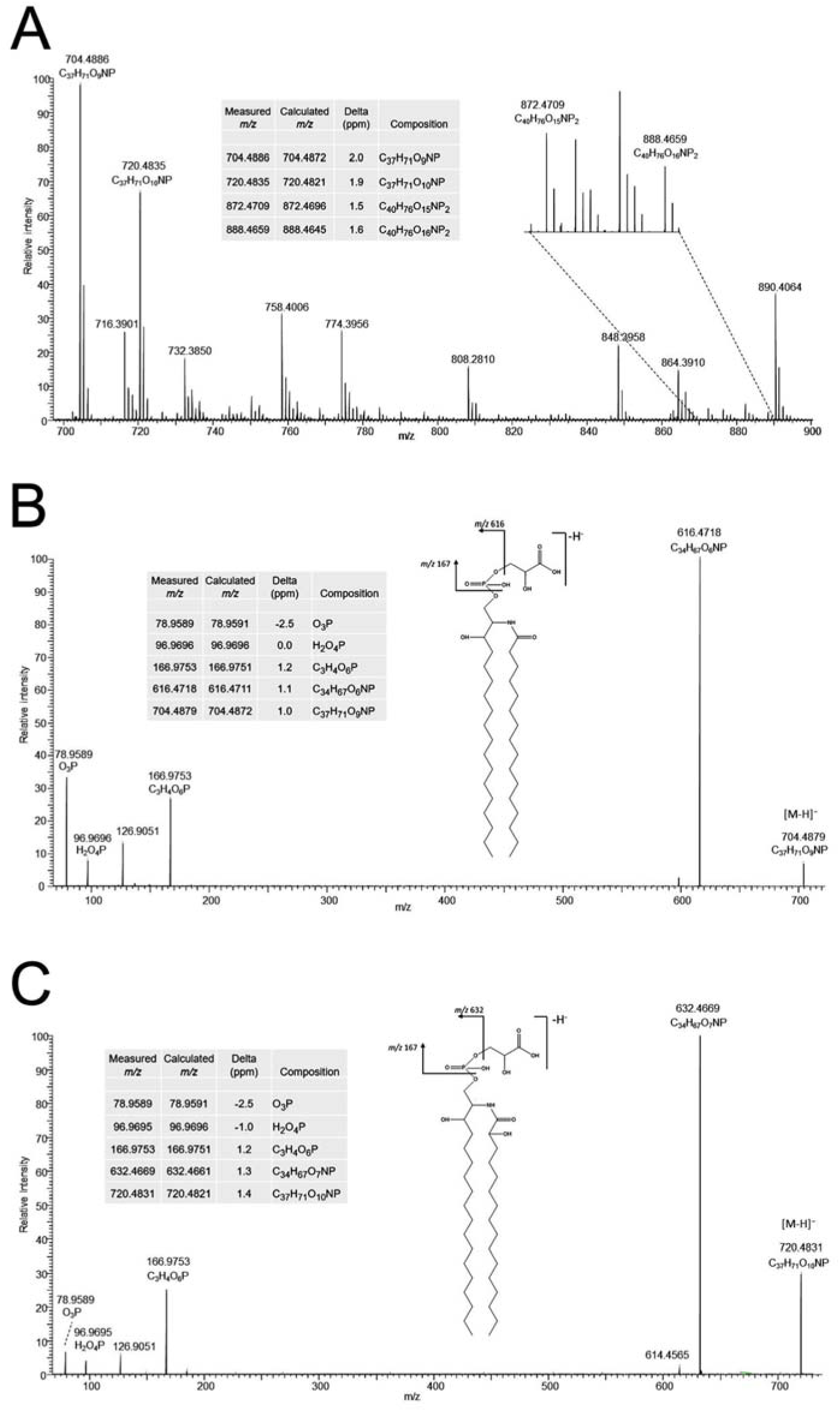
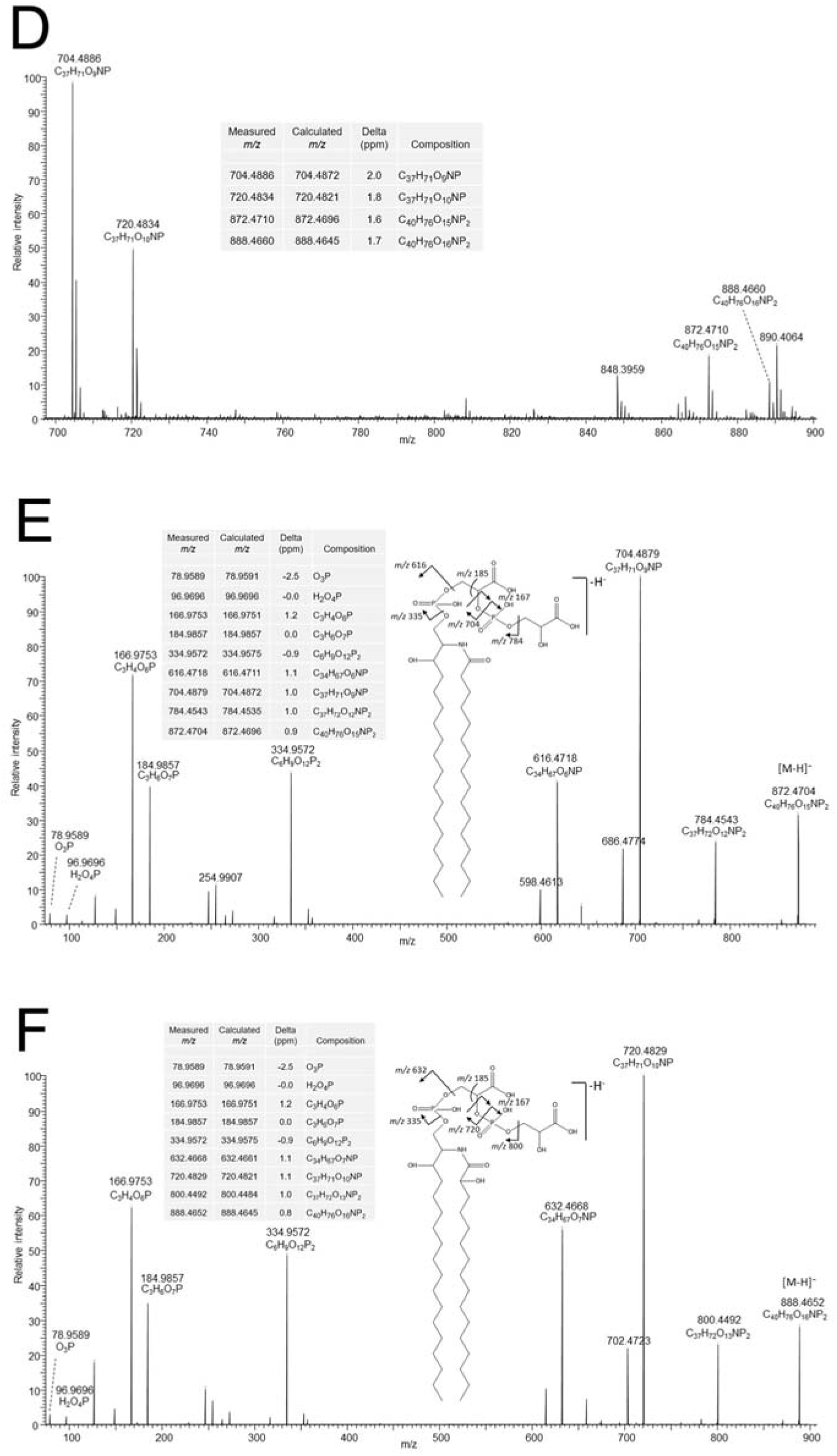
Negative ion mode ESI-Orbitrap mass spectra of lipid extracts of fractions enriched in Lipid I (panel A) and Lipid II (D). Negative ion mode ESI product ion spectra (obtained using HCD with product ions recorded in the Orbitrap) of the M-H^-^ precursor ions at nominal *m/z* 704 (B) and 720 (C) from the fraction enriched in Lipid I, and at *m/z* 872 (E) and 888 (F) from the fraction enriched in Lipid II. Note that the site of fatty acid hydroxylation indicated on the structures is presumed and cannot be assigned to C-2 of the fatty acyl group from the data presented here.

While it is not possible from these mass spectrometric data to determine absolute amounts of the different components, it is possible to consider the relative amounts of the four components in the two extracts. Comparison of the relative signal abundances of the four species in the two extracts makes it clear that the di-phosphoglycerate ceramides are enriched with respect to the mono-phoshoglycerates in the Lipid II-enriched fraction compared with that of Lipid I. We therefore propose that Lipid I is ceramide phosphoglycerate (CPG) and that Lipid II corresponds to ceramide di-phosphoglycerate (CPG2).

### Cofitness and bioinformatic analyses of potential sphingolipid biosynthesis operons in C. crescentus

In our previous work (Olea-Ozuna *et al.,* 2021), we highlighted six operons that are required for high fitness of *C. crescentus* (Christen *et al*., 2011) and that might be involved in SphL biosynthesis and transport (Fig. 4). Specifically, CC_1165, CC_1164, CC_1163, CC_1162, and CC_1154 were shown to be required for the formation of dihydroceramide, the lipidic anchor of SphL (Olea-Ozuna *et al.,* 2021). Notably, other genes in that genomic region of *C. crescentus*, were not required for dihydroceramide synthesis (Olea-Ozuna *et al.,* 2021). However, many mutants affected in genes from this neighborhood show remarkably high cofitness indices with a CC_1162 (*spt*)-deficient mutant (Olea-Ozuna *et al.,* 2021) (Table S1). High cofitness indices (values of 1 or slightly less) between mutants indicate that they display similarly altered phenotypes when compared with wild type, and this suggests that the defective genes might be required for a common specific biochemical pathway (Price *et al.,* 2018). The *spt*-deficient mutant shows moderate cofitness indices (in parentheses) with mutants in CC_1168 (0.52) CC_1167 (0.57), CC_1160 (0.67), CC_1159 (0.60), and CC_1158 (0.60) (Table S1). Another group of genes related by cofitness of their mutants is associated with the predicted 2-phosphoglycerate transferase CC_1159 (Zik *et al*., 2022) with cofitness indices (in parentheses) with mutants in CC_1168 (0.78), CC_1167 (0.54), CC_1161 (0.79), CC_1160 (0.75), CC_1158 (0.72), CC_1156 (0.60), CC_1152 (0.74), and CC_1385 (0.53) (Table S1).

**Fig. 4.**
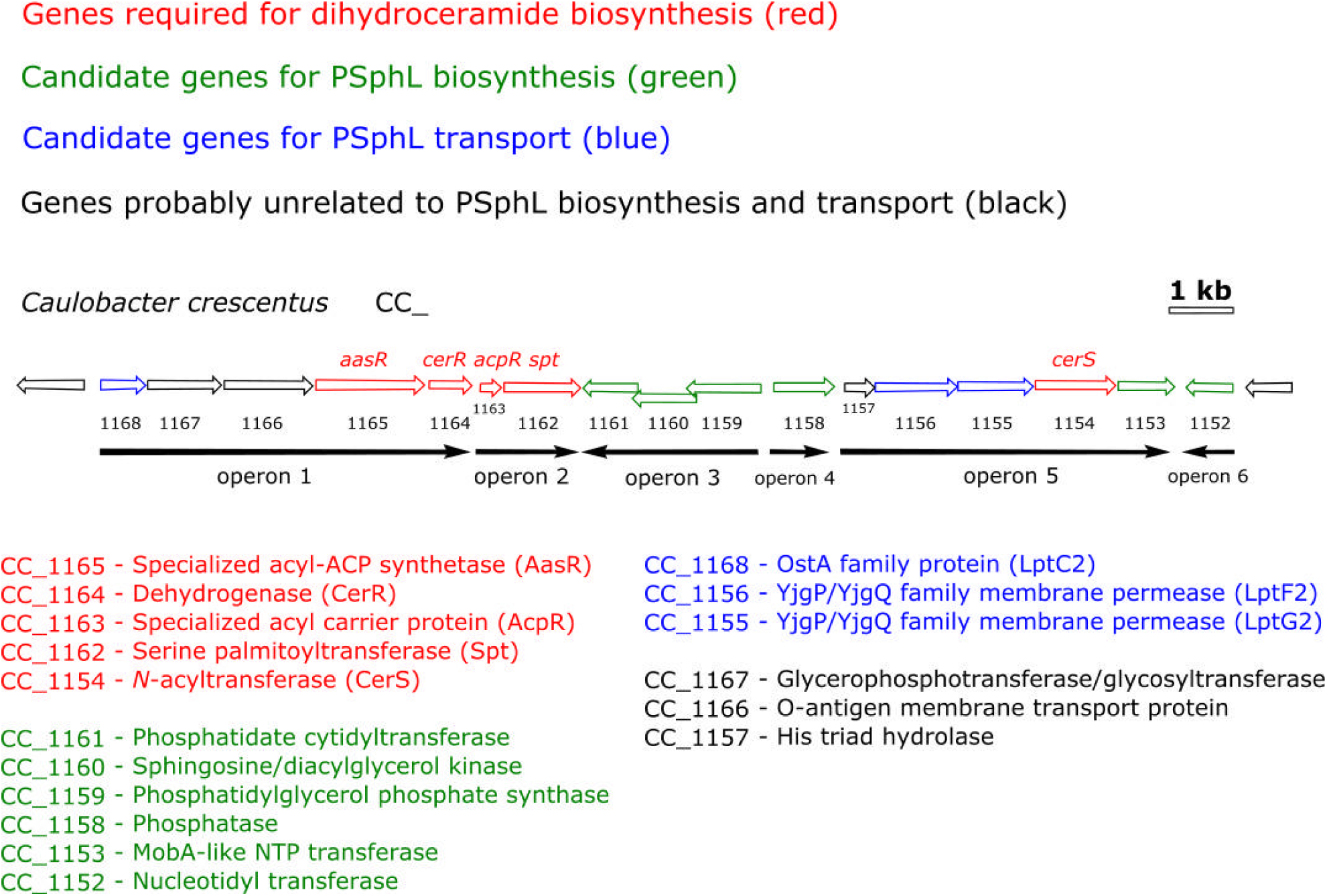
Genomic region with genes of *C. crescentus* involved in the biosynthesis and transport of SphLs. The 17 genes (CC_1168 - CC_1152) of this region are organized into 6 operons and include genes required for dihydroceramide synthesis (CC_1165 - CC_1162, and CC_1154) (Olea-Ozuna *et al*., 2021), potential genes required to convert ceramide to complex PSphLs (CC_1161 - CC_1158, CC_1153, and CC_1152), genes (CC_1168, CC_1156, CC_1155) that code for second versions of LptC, LptF, LptG (i.e. LptC2, LptF2, LptG2) and which probably participate in the selective transport of PSphLs from the IM to the OM, and structural genes probably unrelated to PSphL biosynthesis and transport (CC_1167, CC_1166, and CC_1157). This Figure was modified from Olea-Ozuna *et al*. (2021).

In order to identify candidate genes that might be involved in the conversion of ceramide to PSphLs, we analyzed the genomic region comprising CC_1168 - CC_1152 (Supplementary Results) because many of those genes were linked to ceramide-producing genes through the cofitness results of their respective mutants (Table S1). As a result of these analyses, we suggest that homologs that encode transport proteins (CC_1168, CC_1156, CC_1155) (Fig. 4) might contribute to forming a complex transport system that moves PSphLs from their site of synthesis in the IM to their final destination in the outer layer of the OM in an analogous way to that in which LPS is transported. However, we did not investigate PSphL transport in more detail in this work. Instead, we focused on studying genes potentially coding for biosynthetic enzymes (CC_1161, CC_1160, CC_1159, CC_1158, CC_1153, CC_1152) (Fig. 4) that participate in PSphL biosynthesis.

### CC_1161, CC_1160, CC_1159, CC_1153, and CC_1152 required for the biosynthesis of phospho-sphingolipids

Spt is required for the formation of ceramide, GSphLs (Stankeviciute *et al*., 2019; Olea-Ozuna *et al.,* 2021), and PSphLs (Fig. 1). Separation of ^14^C-acetate-labeled lipids from *C. crescentus* mutants deficient in CC_1161 or CC_1159/CC_1160 produced neither CPG nor CPG2. In contrast, mutants deficient in CC_1153 or CC_1152 produced more intense spots for CPG than wild type but did not produce detectable amounts of CPG2 (Fig. 5A). A mutant deficient in CC_1158 produced slightly less intense CPG2 spots and slightly more intense CPG spots than the wild type (Fig. 5A). When the same strains were radiolabeled with ^32^P-phosphate, results obtained with TLC-separated lipid samples support the previous statements that mutants deficient in CC_1161, CC_1159/CC_1160, CC_1153, or CC_1152 do not form CPG2 (Fig. 5B), that mutants deficient in CC_1153, or CC_1152 displayed elevated levels of CPG, whereas mutants deficient in CC_1161 or CC_1159/CC_1160 were unable to produce CPG. Again, the mutant deficient in CC_1158 produced slightly less intense CPG2 spots and slightly more intense CPG spots than the wild type (Fig. 5B). The accumulation of CPG in mutants deficient in CC_1152, CC_1153, or CC_1158 suggests that these three genes participate in the consumption of CPG and its conversion to CPG2.

**Fig. 5.**
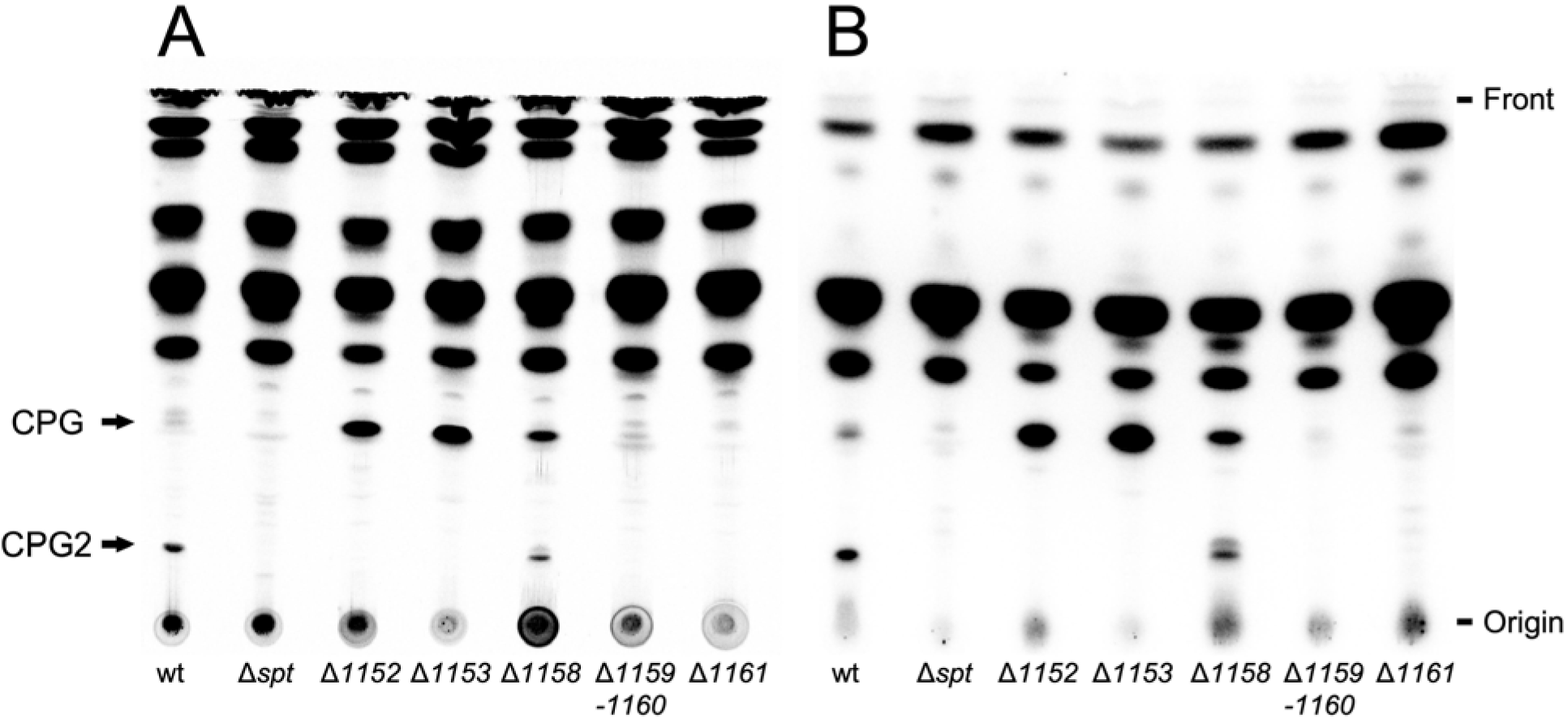
Profile of CPG and CPG2 in mutants deficient in genes required for their formation. Different strains of *C. crescentus* [wild type strain (wt), *spt*-deficient mutant (Δ*spt*), mutant SPG14 deficient in CC_1152 (Δ*1152*), mutant SPG15 deficient in CC_1153 (Δ*1153*), mutant SPG11 deficient in CC_1158 (Δ*1158*), mutant SPG09 deficient in CC_1159/CC_1160 (Δ*1159*/*1160*), and mutant SPG18 deficient in CC_1161 (Δ*1161*)] were cultured in complex medium in the presence of ^14^C-acetate (A) or ^32^P-phosphate (B). After harvesting cells, lipids were extracted, lipid samples were separated by TLC in chloroform/methanol/acetic acid/water (8:3:2:1) and developed chromatograms were analyzed by phosphorimaging. Arrows indicate PSphLs (CPG and CPG2) formed by *C. crescentus*.

Previously we had noted that in a mutant deficient in CC_1159/CC_1160 ceramide levels were much increased when compared to the wild type (Olea-Ozuna *et al*., 2021) and ceramide accumulation in this mutant is now observed again (Fig. S3). Also, in the mutant deficient in CC_1153 much increased ceramide levels were detected, whereas the mutant deficient in CC_1161 displays moderately increased ceramide levels. In mutants deficient in CC_1152 or CC_1158 ceramide levels are below the levels encountered in the wild type (Fig. S3). The accumulation of ceramide in mutants deficient in CC_1153, CC_1161, or CC_1159/CC_1160 suggests that these genes contribute to the consumption of ceramide and its conversion to CPG and CPG2.

When performing complementation experiments, the intact gene(s) were expressed *in trans* in the respective mutants or the mutants harbored an empty vector. Expression of CC_1152 in a CC_1152-deficient mutant shows reduced levels of CPG and increased levels of CPG2 when compared to the CC_1152-deficient mutant harboring the empty vector (Fig. S4A). Also, expression of CC_1153 in a CC_1153-deficient mutant shows reduced levels of CPG and increased levels of CPG2 when compared to the CC_1153-deficient mutant harboring the empty vector. These results suggest that CC_1152 and CC_1153 contribute to the reduction of CPG and are required for the formation of CPG2.

Expression of CC_1158 in a CC_1158-deficient mutant shows reduced levels of CPG and increased levels of CPG2 when compared to the CC_1158-deficient mutant harboring the empty vector (Fig. S4A). Similarly to the mutant deficient in CC_1159/CC_1160 (Fig. 5), the CC_1161 mutant harboring an empty vector did not produce detectable levels of CPG or CPG2. However, when CC_1160 was expressed in the CC_1159/CC_1160-deficient mutant, CPG was formed which was not the case when CC_1159 was expressed instead (Fig. S4A). When expressing CC_1159 and CC_1160 in the CC_1159/CC_1160-deficient mutant, formation of CPG and CPG2 was restored (Fig. S4A). Finally, expression of CC_1161 in a CC_1161-deficient mutant restored formation of CPG and CPG2 which was not the case when the CC_1161-deficient mutant harbored an empty vector (Fig. S4A). When the same strains were radiolabeled with ^33^P-phosphate, mutants in CC_1158, CC_1153, or CC_1152 harboring the empty vector showed increased levels of CPG when compared to the analogous mutants expressing the intact genes *in trans* (Fig. S4B). Expression of CC_1159 in mutants CC_1159/1160 displayed the same lipid profile as this mutant harboring the empty vector (Fig. S4B). Expression of CC_1159 and CC_1160 was required for the formation of CPG2 (Fig. S4B). Notably, expression of CC_1160 in mutant CC_1159/1160 led to increased CPG formation (Fig. S4B). This result indicates that CC_1160, but not CC_1159, is required for CPG formation. When the mutant deficient in CC_1161 was complemented with CC_1161 *in trans*, CPG and CPG2 were formed, which was not the case for the mutant harboring the empty vector (Fig. S4B).

Therefore, TLC studies of lipids of the wild type strain, different mutant strains, and mutant strains expressing intact genes *in trans* in *C. crescentus* suggest that the CC_1161 and CC_1160 genes are required for efficient CPG formation, while the CC_1159, CC_1158, CC_1153, and CC_1152 genes are necessary for the efficient formation of CPG2 (Fig. 6). Although it is tentative to speculate that the ceramide kinase CC_1160 (Zik *et al*., 2022; Dhakephalkar *et al*., 2023) is responsible for the first step and CC_1161 for a subsequent step during CPG formation, we presently do not know in which order the remaining gene products participate in CPG2 formation (Fig. 6). As CC_1152 and CC_1153 are essential for CPG2 formation and as CC_1158 contributes to it, the scheme we propose for CPG2 synthesis is more complex (Fig. 6), than that suggested previously by Zik *et al*. (2022). However, future work needs to clarify the enzymatic functions of CC_1161, CC_1159, CC_1158, CC_1153, and CC_1152 in order to be able to define the precise biosynthesis pathway for CPG and CPG2.

**Fig. 6.**
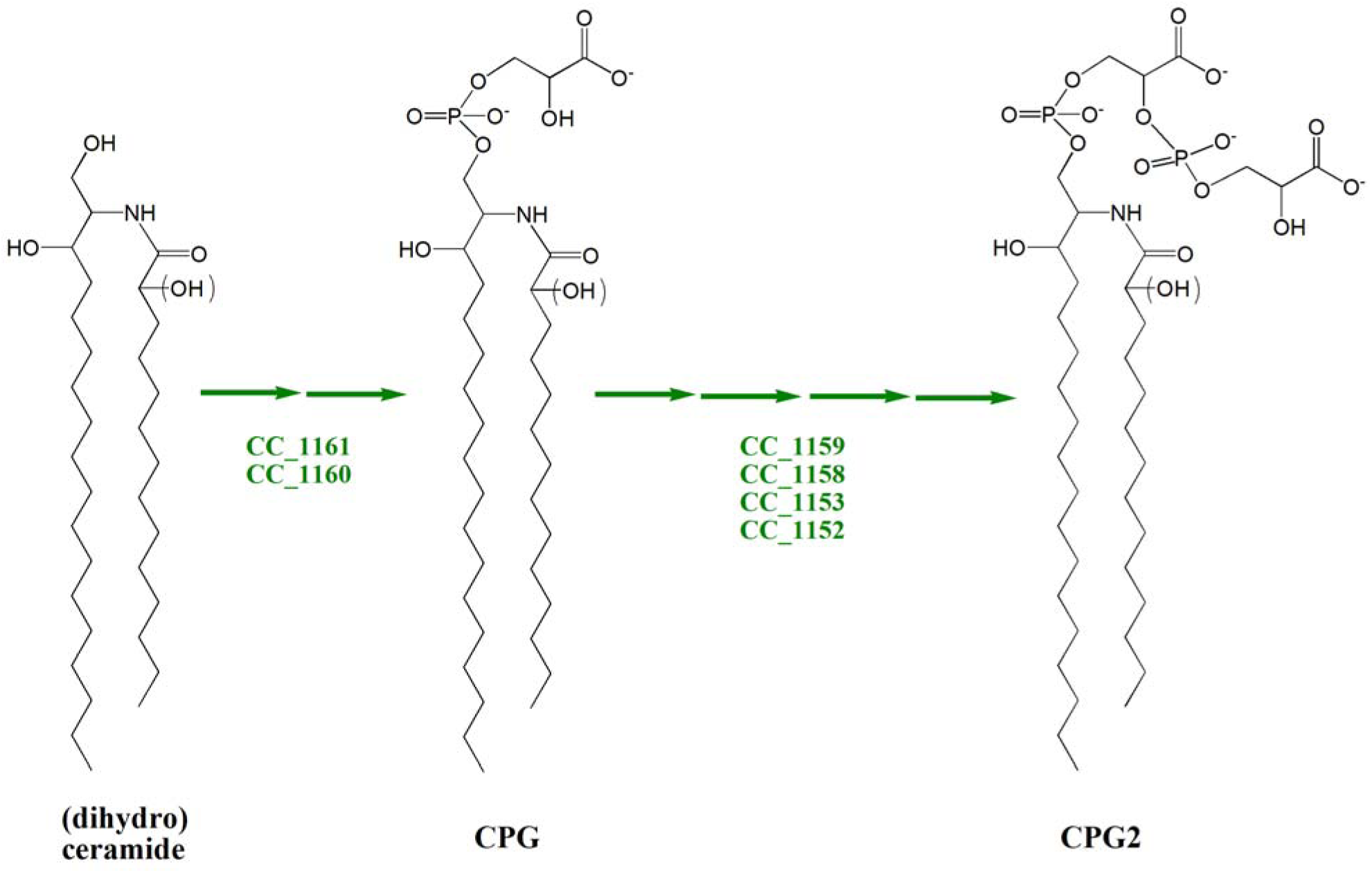
Model for genes involved in biosynthesis of PSphLs CPG and CPG2 in *C. crescentus*. TLC lipid analyses of the wild type strain and the different mutants required in PSphL formation support the model that genes CC_1161 and CC_1160 are required for CPG (Lipid I) formation, while genes CC_1159, CC_1158, CC_1153, and CC_1152 are involved in CPG2 (Lipid II) formation. All three lipid structures shown may carry or not a hydroxy group at an undefined position of their fatty acyl residue.

### Resistance and sensitivity of C. crescentus mutants affected in PSphL biosynthesis towards polymyxin B

In our previous work (Olea-Ozuna *et al*., 2021) we showed that CC_1165, CC_1164, CC_1163, CC_1162, and CC_1154, the genes required for ceramide formation, were needed to confer polymyxin B sensitivity on *C. crescentus*. In the presence of polymyxin B, mutants deficient in CC_1159/CC_1160, CC_1161, or CC_1152 grew much more rapidly than the wild type and similarly to the CC_1162 (*spt*)-deficient mutant (Fig. 7). Expression of CC_1159, CC_1160, or CC_1159/CC_1160 in the CC_1159/CC_1160-deficient mutant reveals that only the combined expression of CC_1159 and CC_1160 restored polymyxin B sensitivity (Fig. S5D). Expression of CC_1161 in the CC_1161-deficient mutant (Fig. S5E), or expression of CC_1152 in a CC_1152-deficient mutant (Fig. S5A), restored polymyxin B sensitivity, whereas the presence of the empty vectors did not. A mutant deficient in CC_1158 grew more rapidly than the wild type, but more slowly than the CC_1162 (*spt*)-deficient mutant in the presence of polymyxin B (Fig. 7), displaying an intermediate phenotype. The expression of intact CC_1158 in the CC_1158-deficient mutant eliminated growth of *C. crescentus* in polymyxin-containing medium (Fig. S5C). Although CC_1153 was thought to be an essential gene (Christen *et al*., 2011), we were able to generate a deletion mutant deficient in CC_1153. However, a CC_1153-deficient mutant grows much more slowly than the wild type in complex PYE medium (Fig. 7A) and this growth behavior is not altered upon cultivation in the presence of polymyxin B (Fig. 7B); therefore, we conclude this mutant is also polymyxin-resistant. Expression of intact CC_1153 in the CC_1153-deficient mutant eliminated growth in polymyxin-containing medium (Fig. S5B), suggesting that CC_1153 contributes to the formation of a PSphL that confers polymyxin sensitivity on *C. crescentus*. Therefore, in addition to the five ceramide biosynthesis genes described previously (Olea-Ozuna *et al*., 2021; Stankeviciute *et al*., 2022; Padilla-Gómez *et al*., 2022), the six genes participating in the conversion of ceramide to the PSphL CPG2 are also required for generating polymyxin sensitivity. LPS of *C. crescentus* lacks phosphate groups and therefore is probably not a target for polymyxin B. In contrast, the negatively charged PSphL CPG2, or a derivative of it, might constitute the main target molecule for polymyxin in *C. crescentus*.

**Fig. 7.**
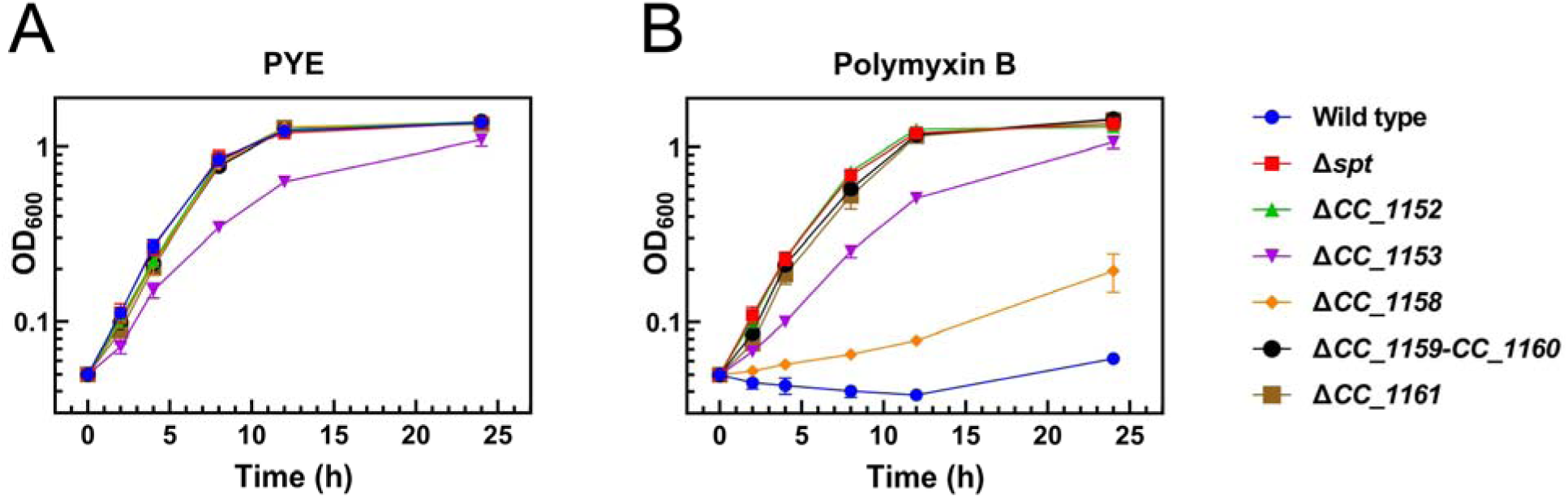
Genes for PSphL formation are required for sensitivity of *C. crescentus* to polymyxin B. Growth (OD_600_) of different strains of *C. crescentus* was determined at 30°C on complex medium (A) or complex medium in the presence of polymyxin B (10 μg/ml) (B). Data and bars represent the average and standard errors obtained from at least three independent experiments.

### Phospho-sphingolipids of C. crescentus are required for resistance towards deoxycholate

The formation of ceramide in *C. crescentus* is required for resistance towards the detergent deoxycholate (Olea-Ozuna *et al*., 2021). We have now studied whether the formation of PSphLs is also required for this resistance. Whereas the wild type strain and the CC_1158-deficient mutant grew well in deoxycholate-containing medium, mutants deficient in CC_1152, CC_1153, CC_1159/CC_1160, and CC_1161, like the *spt*-deficient mutant, did not grow in the presence of deoxycholate (Fig. 8). However, when mutants in CC_1152, CC_1153, CC_1159/CC_1160, and CC_1161 were complemented with the respective intact genes *in trans* they were able to grow in deoxycholate-containing medium, which was not the case if they harbored an empty vector instead (Fig. S6). From these data we conclude that the formation of PSphL CPG2 in *C. crescentus* is required for conferring resistance towards deoxycholate. Therefore, in addition to the five ceramide biosynthesis genes described previously (Olea-Ozuna *et al*., 2021; Stankeviciute *et al*., 2022; Padilla-Gómez *et al*., 2022), five genes participating in the conversion of ceramide to CPG2 are also required for generating resistance towards the detergent deoxycholate and therefore for membrane stability. The fact that the mutant deficient in CC_1158 grows like the wild type in the presence of deoxycholate suggests that in this mutant enough CPG2 or a derivative of it is still formed to provide deoxycholate resistance.

**Fig. 8.**
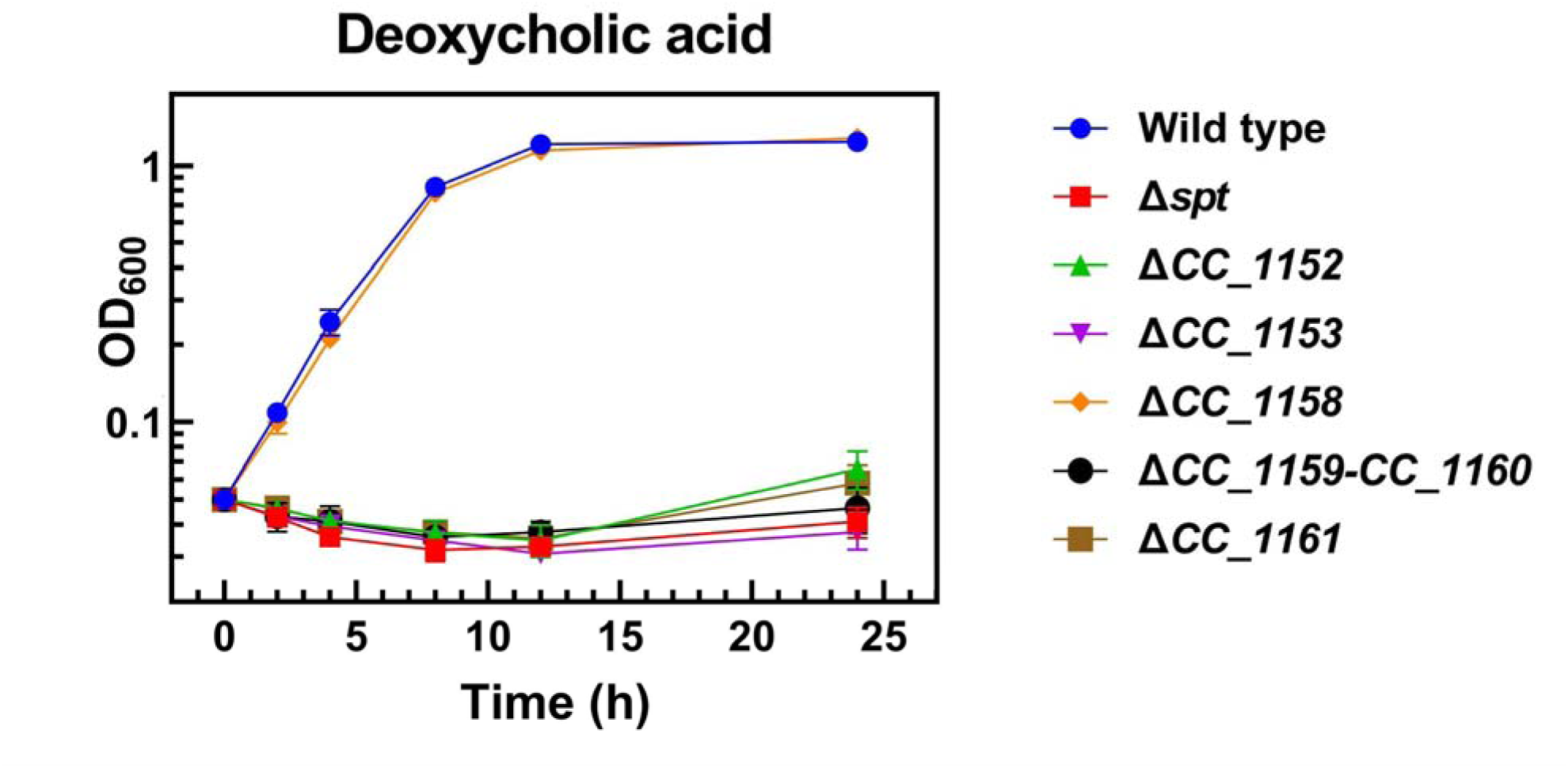
Genes for PSphL formation are required for resistance of *C. crescentus* to deoxycholate. Growth (OD_600_) of different strains of *C. crescentus* [wild type strain (Wild type), *spt*-deficient mutant (Δ*spt*), mutant SPG14 deficient in CC_1152 (Δ*1152*), mutant SPG15 deficient in CC_1153 (Δ*1153*), mutant SPG11 deficient in CC_1158 (*Δ1158*), mutant SPG09 deficient in CC_1159/CC_1160 (Δ*1159*-*1160*), and mutant SPG18 deficient in CC_1161 (Δ*1161*)] was determined at 30°C on complex medium in the presence of deoxycholate (1 mg/ml). Data and bars represent the average and standard errors obtained from at least three independent experiments.

### Phospho-sphingolipids of C. crescentus are required for virulence on Galleria mellonella larvae

In recent years, larvae of the greater wax moth *Galleria mellonella* have become an important animal model for studying virulence. Pathogens injected into *G. mellonella* provoke cellular and humoral responses by the larvae (Pereira *et al*., 2018). Notable humoral immune responses include the formation of antimicrobial peptides and the activation of phenol oxidase leading to melanin formation and melanization of the larvae when counteracting an immune challenge due to invasive microbes (Pereira *et al*., 2018). Recent studies have shown that distinct *Caulobacter* species are virulent in the *G. mellonella* infection model (Moore and Gitai, 2020). In order to analyze whether bacterial SphLs might be responsible for this virulence, we tested the effect of various *C. crescentus* strains when injected into *G. mellonella* larvae. Over a period of five days, we distinguished healthy larvae from melanized and dead larvae.

When setting up healthspan assays, we initially inoculated PYE medium, nonvirulent *E. coli*, or virulent *Burkholderia cenocepacia* (Córdoba-Castro *et al*., 2021) strains into *G. mellonella* larvae. Injection of PYE medium or of *E*. *coli* did not decrease the percentage of healthy larvae in a significant way over a 5 day period (Fig. S7). In contrast, injection of *B*. *cenocepacia* led to a steady decline in healthy larvae over the first 3 days (Fig. S7); by then all larvae were either melanized or dead.

Whereas healthy larvae declined rapidly when inoculated with *C. crescentus* wild type, this was not the case when an *spt*-deficient mutant of *C. crescentus* was applied instead (Fig. 9AB). An *spt*-deficient mutant complemented with the intact *spt* gene *in trans* caused rapid decrease of healthy larvae and therefore restored *C. crescentus* virulence in *G. mellonella,* whereas the *spt*-deficient mutant harboring an empty vector did not (Fig. 9A). Therefore, as Spt is required for the formation of any SphL, bacterial SphLs are required for *C. crescentus* virulence. Remarkably, most of the melanization provoked by *C. crescentus* in *G. mellonella* larvae occurred within the first 24 h after injection (Fig. 9AB), but even after 5 days a few larvae had survived and were not melanized. Statistical analyses show that the percentage of melanized larvae after 24 h is significantly superior when inoculated with *C. crescentus* wild type than with the *spt*-deficient mutant (Fig. 9B). Also, inoculation of larvae with a *spt*-deficient mutant complemented with the intact *spt* gene *in trans* caused a significantly higher percentage of melanized larvae when compared with larvae that had been inoculated with a *spt*-deficient mutant harboring an empty vector (Fig. 9B).

**Fig. 9.**
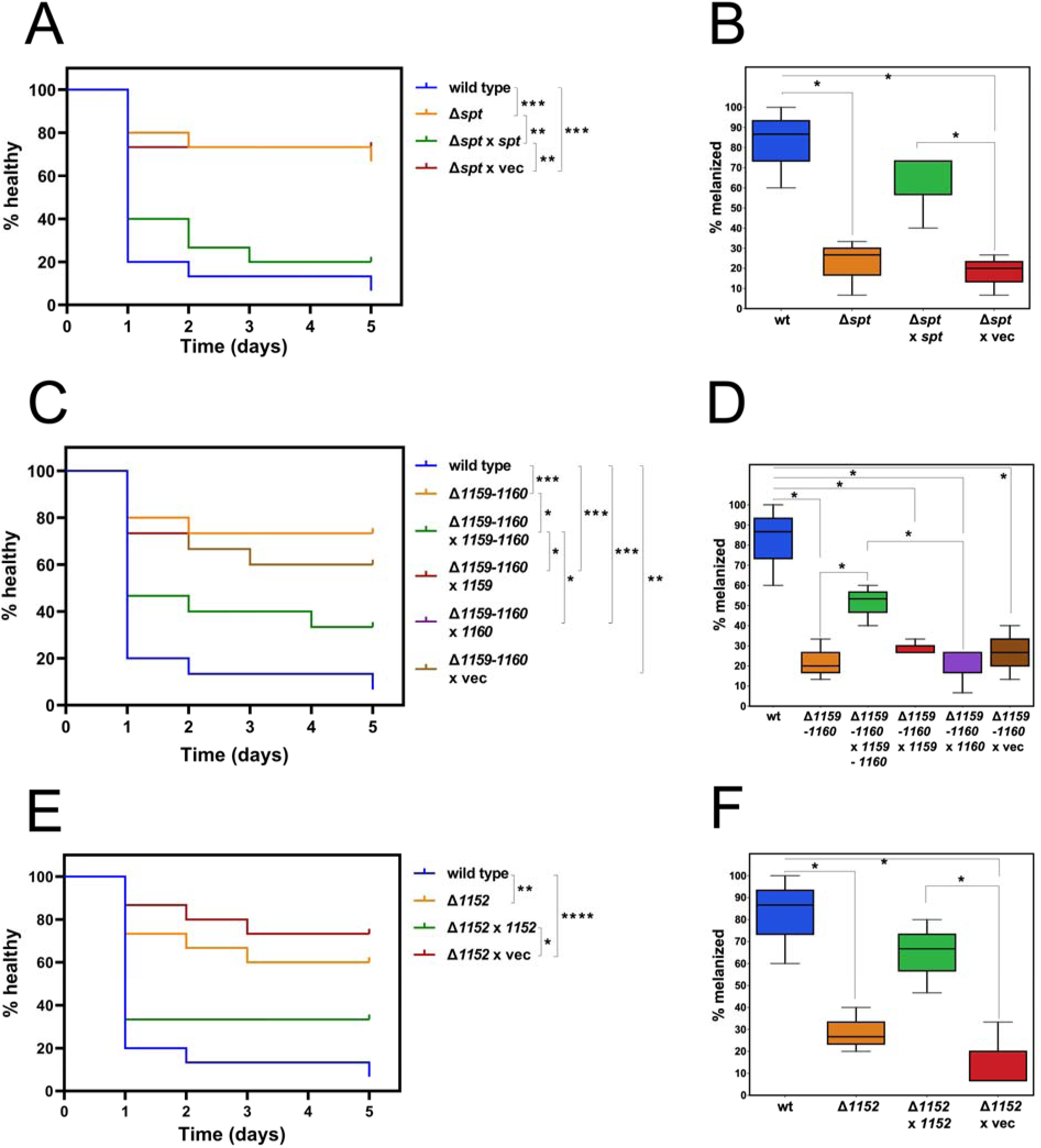
PSphL formation is required for virulence of *C. crescentus* in *G. mellonella* larvae. Healthspan of *G. mellonella* larvae was determined with Kaplan-Meier survival analysis up to 5 days after inoculation with distinct *C. crescentus* strains (left column). Melanized *G. mellonella* larvae 24 h after inoculation of the larvae with distinct *C. crescentus* strains (right column). Inoculation of *G. mellonella* larvae was performed with 10^5^ CFU of wild type (wt) *C. crescentus* strain, *spt*-deficient mutant (Δ*spt*), *spt*-deficient mutant expressing intact *spt in trans* (Δ*spt* x *spt*), *spt-*deficient mutant harboring the empty vector (Δ*spt* x vec)(A, B), CC_1159/CC_1160-deficient mutant (Δ*1159-1160*), CC_1159/CC_1160-deficient mutant expressing intact CC_1159/CC_1160 *in trans* (Δ*1159-1160* x *1159-1160*), CC_1159/CC_1160-deficient mutant expressing intact CC_1159 *in trans* (Δ*1159-1160* x *1159*), CC_1159/CC_1160-deficient mutant expressing intact CC_1160 *in trans* (Δ*1159-1160* x *1160*), CC_1159/CC_1160-deficient mutant harboring the empty vector (Δ*1159-1160* x vec)(C, D), CC_1152-deficient mutant (Δ*1152*), CC_1152-deficient mutant expressing intact CC_1152 *in trans* (Δ*1152* x *1152*), or CC_1152-deficient mutant harboring the empty vector (Δ*1152* x vec)(E, F), respectively. Survival curves are a representative cohort (n = 15) of at least three biological replicates (Mantel-Cox test for statistics, *P<0.5, **P<0.01, ***P<0.001, ****P<0.0001).

Whereas healthy larvae declined rapidly when inoculated with *C. crescentus* wild type, this was not the case when a CC_1159/CC_1160-deficient mutant of *C. crescentus* was injected instead (Fig. 9CD). A CC_1159/CC_1160-deficient mutant complemented with the intact CC_1159/CC_1160 genes caused a rapid decrease in healthy larvae and therefore restored virulence in *G. mellonella* whereas the CC_1159/CC_1160-deficient mutant harboring an empty vector *in trans* did not (Fig. 9CD). Expression of CC_1159 or CC_1160 alone in the CC_1159/CC_1160-deficient mutant did not restore virulence of *C. crescentus* in *G. mellonella* larvae (Fig. 9CD) either. Statistical analyses show that the percentage of melanized larvae after 24 h is significantly superior when inoculated with *C. crescentus* wild type than with the CC_1159/CC_1160-deficient mutant (Fig. 9D). In addition, inoculation of larvae with a CC_1159/CC_1160-deficient mutant complemented with the intact CC_1159/CC_1160 genes *in trans* caused a significantly higher percentage of melanized larvae when compared with larvae that had been inoculated with a CC_1159/CC_1160-deficient mutant harboring an empty vector, or vectors expressing only the CC_1159 or CC_1160 gene (Fig. 9D). As CC_1160 is required for the formation of any PSphL (Figs. 5 and S4), bacterial PSphLs are required for *C. crescentus* virulence.

Similarly, whereas healthy larvae declined rapidly when inoculated with *C. crescentus* wild type, this was not the case when a CC_1152-deficient mutant of *C. crescentus* was injected instead (Fig. 9EF). A CC_1152-deficient mutant complemented with the intact CC_1152 gene caused a rapid decrease in healthy larvae and therefore restored virulence in *G. mellonella* whereas the CC_1152-deficient mutant harboring an empty vector *in trans* did not (Fig. 9EF). Statistical analyses show that the percentage of melanized larvae after 24 h is significantly superior when inoculated with *C. crescentus* wild type than with the CC_1152-deficient mutant (Fig. 9F). Inoculation of larvae with a CC_1152-deficient mutant complemented with the intact CC_1152 gene *in trans* caused a significantly higher percentage of melanized larvae when compared with larvae that had been inoculated with a CC_1152-deficient mutant harboring an empty vector (Fig. 9F). As CC_1152 is required for the formation of CPG2, this bacterial PSphL or derivatives of it are required for *C. crescentus* virulence.

In previous work, Moore and Gitai (Moore and Gitai, 2020) suggested that the virulence-provoking compound was contained in the LPS fraction. However, this present study shows that genes required for SphL, PSphL, and specifically CPG2 formation are needed for *C. crescentus* virulence on larvae of the greater wax moth *G. mellonella*.

## Conclusion

The present analysis suggests that besides the five genes required for ceramide synthesis in *C. crescentus*, another six genes (CC_1161, CC_1160, CC_1159, CC_1158, CC_1153, CC_1152) from this genomic region participate in the formation of the PSphL CPG2, the presumptive target molecule for interaction with polymyxin, and therefore responsible for the trait of polymyxin sensitivity in *C*. *crescentus*. Ten of the eleven genes required for the biosynthesis of the PSphL CPG2 are needed for an intact bacterial cell envelope and for the formation of a cell-associated factor that provokes virulence on larvae of the greater wax moth *G*. *mellonella*. The mutant deficient in CC_1158 often shows intermediate phenotypes, which might suggest the existence of another gene that can partially substitute for CC_1158.

## Experimental Procedures

### Bacterial strains, plasmids, and growth conditions

The bacterial strains and plasmids used and their relevant characteristics are shown in Table 1. The construction of caulobacterial mutants deficient in putative SphL biosynthesis genes is described in Table S2. Strains of *C. crescentus, Escherichia coli* MG1655, or *Burkholderia cenocepacia* J2315 were grown in complex peptone yeast extract (PYE) medium (Ely, 1991) at 30°C on a gyratory shaker at 250 rpm. For growth experiments, strains were first grown on PYE plates. Then, cells were resuspended at cell densities of 5 x 10^7^ cells/ml in liquid PYE medium and grown for 20 h during such a first growth cycle. During a second subcultivation in PYE medium, again inoculating with 5 x 10^7^ cells/ml, growth of *C. crescentus* wild type and of SphL-deficient mutants was followed by determining OD_600_. For studying bacterial growth in the presence or absence of polymyxin B, deoxycholate, cultures with 15 ml medium in 125 ml Erlenmeyer flasks were employed.

**Table 1.**
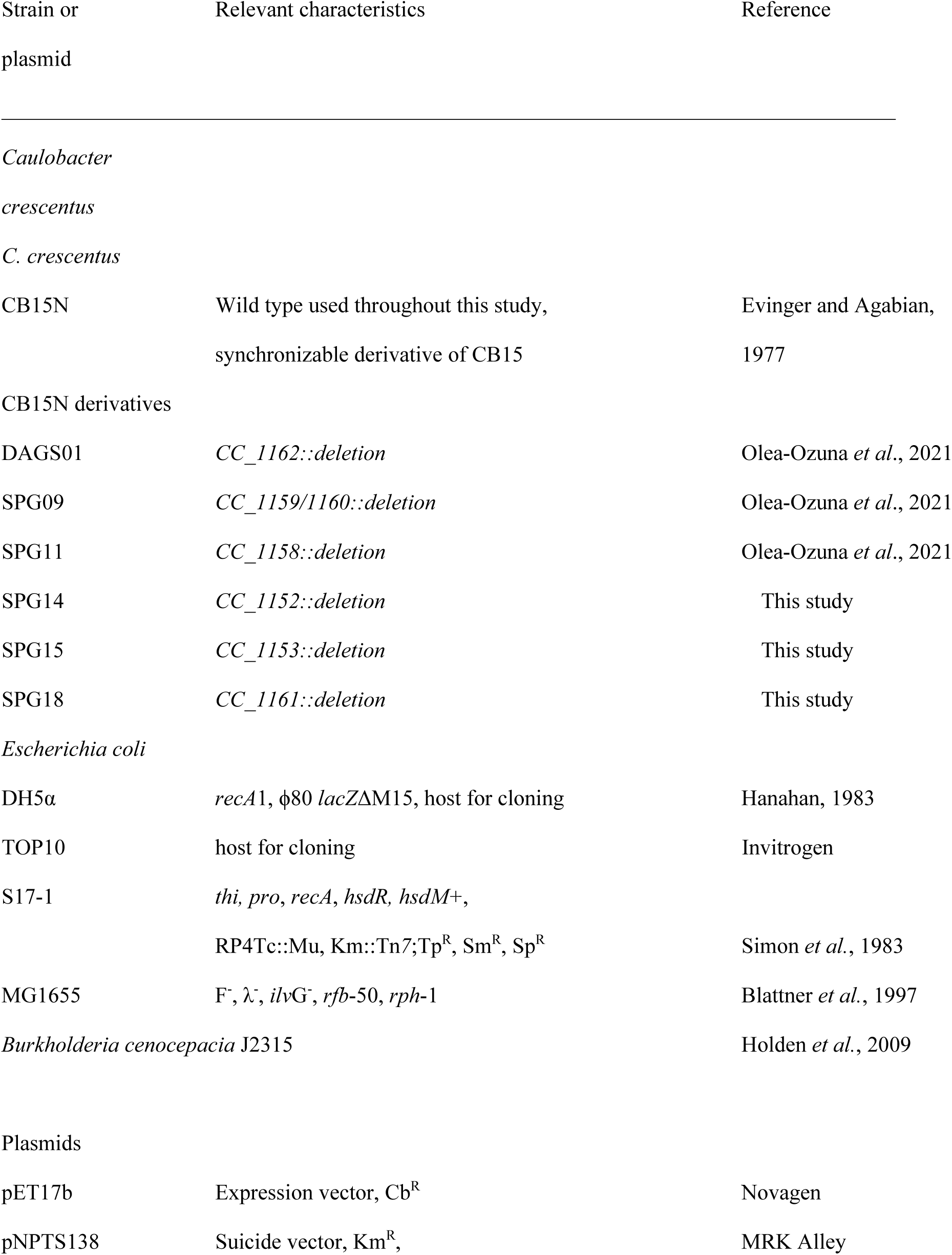
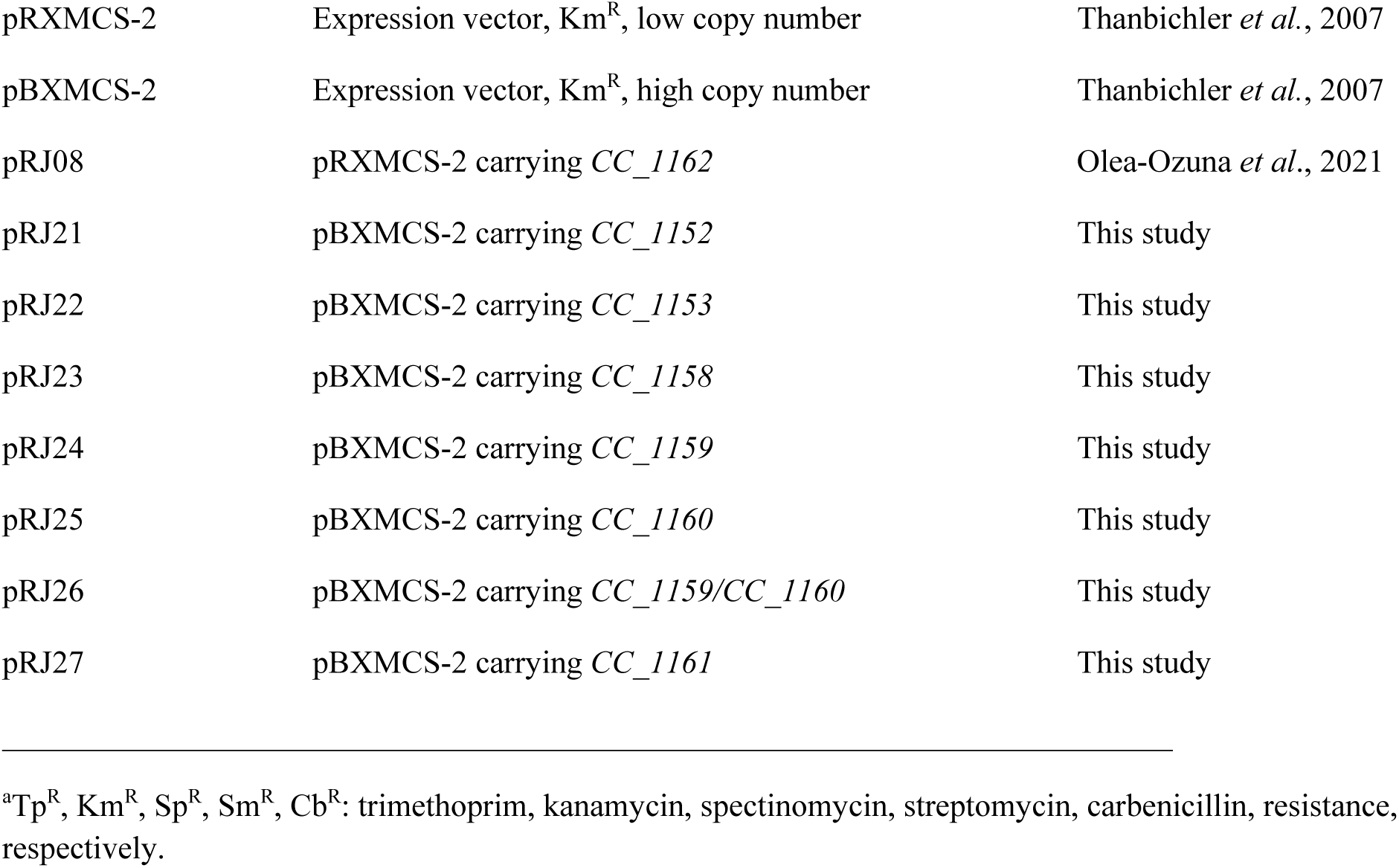
Bacterial strains and plasmids.

Other *E. coli* strains were cultured on Luria-Bertani (LB) medium (Miller, 1972) at 37°C. Antibiotics were added to media in the following concentrations (μg/ml) when required: kanamycin 5 (25 in solid media), polymyxin B 10, in the case of *C. crescentus*, and kanamycin 50, carbenicillin 50, in the case of *E. coli*.

Plasmids pBXMCS-2, pNPTS138, and their derivatives were moved into *C. crescentus* by conjugation. In the conjugation, aliquots of exponentially growing cultures of a donor strain of *E. coli* S17-1, harboring the plasmid of interest (200 μl), and of the receiver strain *C. crescentus* CB15N (1 ml) were mixed. The mixed cell suspension was centrifuged and washed three times with ice-cold 10% glycerol, resuspended in 100 μl of PYE medium and applied in drops onto PYE agar. After most of the liquid had evaporated, the cell mixture was incubated at 30°C for 16 h and potential transconjugants were selected on PYE agar containing nalidixic acid (25 μg/ml) and kanamycin (25 μg/ml).

### DNA manipulations and analyses

Recombinant DNA techniques were performed according to standard protocols (Sambrook *et al*., 2001). Commercial sequencing of amplified genes by Eurofins Medigenomix (Martinsried, Germany) corroborated the correct DNA sequences. The DNA regions containing *cc_1168-cc_1152* were analyzed using the NCBI (National Center for Biotechnology Information) BLAST network server (Altschul *et al*., 1997).

### Construction of expression plasmids

Using PCR and a pair of specific oligonucleotides (Table S3) genes or combinations of genes encoding potential structural genes for SphL biosynthesis were amplified from *C. crescentus* genomic DNA. Suitable restriction sites for cloning of the genes were introduced by PCR with oligonucleotides. After restriction with the respective enzymes, the PCR-amplified DNA fragments were cloned into a pET17b or a pBXMCS-2 or recloned into a pBXMCS-2 vector as detailed in Table S3.

### In vivo labeling of bacterial strains with [^14^C] acetate, [^32^P] phosphate or [^33^P] phosphate

The lipid composition of bacterial strains was determined after labeling with [1-^14^C] acetate (60 mCi/mmol; Perkin Elmer), [^32^P] phosphate (1 Ci/mmol; Perkin Elmer), or [^33^P] phosphate (1 Ci/mmol; American Radiolabeled Chemicals, Inc.). Cultures (1 ml) of wild type or mutant strains were inoculated from precultures grown in the same medium in order to obtain an initial cell density of 2 x 10^8^ cells/ml. After the addition of 1 μCi [1-^14^C] acetate, 2.5 μCi [^32^P] phosphate, or of 2.5 μCi [^33^P] phosphate to each culture, they were incubated for 16 h. At the end of the respective incubation periods, cells were harvested by centrifugation, and resuspended in 100 µl of water. Lipids were extracted according to the method of Bligh and Dyer (1959) and the chloroform phase was separated by one-dimensional TLC on high performance TLC aluminum sheets (silica gel 60; Merck Poole, United Kingdom) and developed with chloroform/methanol/ammonium hydroxide (40:10:1; v/v) or chloroform/methanol/acetic acid/water (8:3:2:1; v/v) as the mobile phase. Radioactive lipids were visualized by phosphorimaging using a Typhoon FLA 9500 and quantification was performed with Image Quant TL (Amersham Biosciences).

### Mass spectrometric analysis

For mass spectrometric analyses, usually bacterial cultures comprising 100 ml of medium were grown, lipids were extracted as described above (Bligh and Dyer, 1959) and studied directly. In the case of enriched Lipid I or Lipid II fractions, the *spt*-deficient mutant expressing *spt in trans* or harboring an empty vector, was cultivated in 2 l of medium to an OD_600_ of 1.2, when cells were harvested, lipids extracted, and separated by TLC on silica gel-containing plates. After separation with chloroform/methanol/acetic acid/water (8:3:2:1; v/v) as the mobile phase, lipids were visualized using iodine vapor and silica gel fractions containing Lipid I or Lipid II were scraped from the plates and lipids were extracted from the silica gels. Aliquots of the enriched Lipid I or Lipid II fractions were reanalyzed by TLC and did not contain major other lipidic compounds that could be detected by staining with iodine vapor (Sims and Larose, 1961) or with 8-anilo-1-naphthalenesulfonic acid (ANS) reagent (Zbierzak *et al*., 2011).

High resolution mass spectra were acquired with a solariX XR FT-ICR mass spectrometer equipped with a 9.4 T superconducting magnet (Bruker Daltonics) and an Orbitrap Fusion Tribrid mass spectrometer (Thermo Scientific). Spectra were acquired in negative ion mode using ESI; lipid samples were redissolved in dichloromethane:methanol (1:1, v:v) and further diluted in methanol; solutions were introduced into the ion source by syringe infusion with a flow rate of 2 - 3 μL min^−1^.

SolariX: ESI conditions were spray voltage 3500 V, nebuliser gas pressure 2 bar, drying gas flow 4 L min^−1^; temperature 160 °C. External calibration was performed on sodium formate clusters with a 10 μg mL^−1^ solution in 50% propan-2-ol. Mass spectra were also internally calibrated on fatty acid peaks using linear calibration. This procedure results in a mass accuracy better than 0.5 ppm for the mass spectrometric signals reported (2 ppm for product ion signals). The spectra were acquired with 1 M data points over the *m/z* range 100–2000 (transient of 0.367 s) resulting in a resolving power of 37000 at *m/z* 700. CID was performed by isolation (width *m/z* 2.0) of the precursor ions in the quadrupole and then storage in the hexapole collision cell with an excitation voltage of 23 V. The ion accumulation time in the ion source was set to 0.1 s (0.5 s for product ion spectra). A total of 8 scans were added for each mass spectrum. Spectra visualization and formula calculation was performed with DataAnalysis 5.0 (Bruker Daltonics).

Orbitrap Fusion: HESI source settings were spray voltage 3300 V, sheath gas 2.5 Arb, aux gas 0.5 Arb, ion transfer tube temperature 275 °C. Spectra were acquired with 120000 resolution at *m/z* 200 (64000 at *m/z* 700), scan range *m/z* 200 – 1000 for mass spectra and auto for product ion spectra. Easy-IC was used for internal calibration. For production spectra, isolation (width *m/z* 1.6) was carried out in the quadrupole followed by HCD with excitation voltage 25 – 32 V. A total of approximately 1 minute of acquired data were averaged to produce each spectrum. Spectra visualization and formula calculation was performed with Xcalibur (version 4.0; Thermo Scientific).

### Galleria mellonella healthspan assay

All *G. mellonella* larvae were obtained from PETMMAL, S.A. DE C.V. (Cuautitlán Izcalli, México) and were kept in an incubator at 30°C. The larvae were used for health tests within three days of receiving the package. For virulence experiments, *C. crescentus* strains were resuspended to an OD_600_ of 0.05 and in PYE medium and cultivated until reaching an OD_600_ of 0.125 (10^7^ CFU/ml). Cells were washed three times with PYE medium and resuspended in the original volume. A 31G (gauge) insulin syringe was used to inoculate 10 μl of bacterial suspension, which corresponds to 10^5^ CFU. Virulence was tested using 3 cohorts of 15 *G. mellonella* larvae placed in Petri dishes without diet and incubated at 30°C. Melanization and mortality were assessed every 24 h for 5 days after injection. Healthy larvae were neither melanized nor dead.

## Acknowledgements

R.J.O.O. was a Ph.D. student from the Programa de Doctorado en Ciencias Biomédicas (PDCB), Universidad Nacional Autónoma de México (UNAM) and received fellowship 621047 from Consejo Nacional de Ciencia y Tecnología de México (CONACyT) (CVU 822843). We thank Angeles Moreno and Miguel Ángel Vences-Guzmán for skillful technical assistance. This work was supported by grants from DGAPA/UNAM (IN201120 and IN202223), CONACyT/Mexico (178359 and 253549 in Investigación Científica Básica as well as 118 in Investigación en Fronteras de la Ciencia). The mass spectrometric analyses were carried out in the York Centre of Excellence in Mass Spectrometry, which was created thanks to a major capital investment through Science City York, supported by Yorkshire Forward with funds from the Northern Way Initiative, and subsequent support from EPSRC (EP/K039660/1; EP/M028127/1). This article is dedicated to the memory of the late Franz Martin Lingens.

## References

Altschul, S.F., Madden, T.L., Schäffer, A.A., Zhang, J., Zhang, Z., Miller, W., and Lipman, D.J. (1997) Gapped BLAST and PSI BLAST: A new generation of protein database search programs. Nucleic Acids Res 25: 3389–3402. doi: 10/1093/nar/25.17.3389

Bertani, B., and Ruiz, N. (2018) Function and Biogenesis of Lipopolysaccharides. EcoSal Plus, 8(1), 10.1128/ecosalplus.ESP-0001-2018.

Blattner, F.R., Plunkett 3^rd^, G., Bloch, C.A., Perna, N.T., Burland, V., Riley, M. et al. (1997) The complete genome sequence of *Escherichia coli* K-12. Science 277: 1453–1462. doi: 10.1026/science.277.5331.1453

Bligh, E.G., and Dyer, W.J. (1959) A rapid method for total lipid extraction and purification. Can J Biochem Physiol 37: 911–917.

Bridger, N., Walkty, A., Crockett, M., Fanella, S., Nichol, K., and Karlowsky, J.A. (2011) *Caulobacter* species as a cause of postneurosurgical bacterial menigitis in a pediatric patient. Can J Infect Dis Med Microbiol 22: e10–e12

Christen, B., Abeliuk, E., Collier, J.M., Kalogeraki, V.S., Passarelli, B., Coller, J.A., et al. (2011) The essential genome of a bacterium. Mol Syst Biol 7:528. doi: 10.1038/msb.2011.58.

Córdoba-Castro, L.A., Salgado-Morales, R., Torres, M., Martínez-Aguilar, L., Lozano, L., Vences-Guzmán, M.A., et al. (2021) Ornithine lipids in *Burkholderia* spp. pathogenicity. Front Mol Biosci 7: 10932. doi: 10.3389/fmolb.2020.610932

Dhakephalkar, T., Stukey, G.J., Guan, Z., Carman, G.M., and Klein, E.R. (2023) Characterization of an evolutionary distinct bacterial ceramide kinase from *Caulobacter crescentus*. J Biol Chem 299: 104894. doi: 10.1016/j.jbc.2023.104894

Doerrler W. T. (2006) Lipid trafficking to the outer membrane of Gram-negative bacteria. Mol Microbiol 60: 542–552.

Ely, B. (1991) Genetics of *Caulobacter crescentus*. Methods Enzymol 204: 372– 384.

Evinger, M., and Agabian, N. (1977) Envelope-associated nucleoid from *Caulobacter crescentus* stalked and swarmer cells. J Bacteriol 132: 294–301.

Geiger, O., González-Silva, N., López-Lara, I.M., and Sohlenkamp, C. (2010) Amino acid-containing membrane lipids in bacteria. Prog Lipid Res 49: 46– 60.

Geiger, O., Padilla-Gómez, J., and López-Lara, I.M. (2019) Bacterial sphingolipids and sulfonolipids. In Handbook of Hydrocarbon and Lipid Microbiology; Biogenesis of Fatty Acids, Lipids and Membranes, O. Geiger (ed). Cham, Switzerland: Springer Nature Switzerland AG, pp. 123–137.

Hanahan, D. (1983) Studies on transformation of *Escherichia coli* with plasmids. J Mol Biol 166: 557–580.

Holden, M.T., Seth-Smith, H.M., Crossman, L.C., Sebaihia, M., Bentley, S.D., Cerdeno-Tarraga, A.M. et al. (2009) The genome of *Burkholderia cenocepacia* J2315, an epidemic pathogen of cystic fibrosis patients. J Bacteriol 191: 261–277. [Corrigendum: *J Bacteriol* (2009) 191: 2907.]

Justesen, U.S., Holt, H.M., Thiesson, H.C., Blom, J., Nielson, X.C., Dargis, R., Kemp, M., and Christensen, J.J. (2007) Report of the first human case of *Caulobacter* sp. infection. J Clin Microbiol 45: 1366–1369.

Kawasaki, S., Moriguchi, R., Sekiya, K., Nakai, T., Ono, E., Kume, K., and Kawahara, K. (1994) The cell envelope structure of the lipopolysaccharide-lacking gram-negative bacterium *Sphingomonas paucimobilis*. J Bacteriol 176: 284– 290.

Keck, M., Gisch, N., Moll, H., Vorhölter, F.J., Gerth, K., Kahmann, U., et al. (2011) Unusual outer membrane lipid composition of the gram-negative, lipopolysaccharide-lacking myxobacterium *Sorangium cellulosum* So ce56. J Biol Chem 286: 12850– 12859.

Miller, J.H. (1972) Experiments in Molecular Genetics. Plainview, NY, USA: Cold Spring Harbor Laboratory Press.

Moore, G.M., and Gitai, Z. (2020) Both clinical and environmental *Caulobacter* species are virulent in the *Galleria mellonella* infection model. PLos ONE 15(3): e0230006. Doi.org/10-1371/journal.pone.0230006

Nelson, D.L., and Cox, M.M. (2021). Lehninger: Principles of Biochemistry, 8th ed. New York, USA: W.H. Freeman and Company.

Nikaido, H. (2003) Molecular basis of bacterial outer membrane permeability revisited. Microbiol Mol Biol Rev 67(4): 593–656.

Okuda, S., Sherman, D. J., Silhavy, T. J., Ruiz, N., and Kahne, D. (2016) Lipopolysaccharide transport and assembly at the outer membrane: the PEZ model. Nature Rev Microbiol 14: 337– 345.

Olea-Ozuna, R.J., Poggio, S., Bergström, E., Quiroz-Rocha, E., García-Soriano, D.A., Sahonero-Canavesi, D.X., Padilla-Gómez, J., Martínez-Aguilar, L., López-Lara, I.M., Thomas-Oates, J., Geiger, O. (2021) Five structural genes required for ceramide synthesis in *Caulobacter* and for bacterial survival. Environ Microbiol 23: 143–159. doi: 10.1111/1462-2920.15280

Padilla-Gómez, J., Olea-Ozuna, R.J., Contreras-Martínez, S., Morales-Tarré, O., García-Soriano, D.A. Sahonero-Canavesi, D.X., Poggio, S., Encarnación-Guevara, S., López-Lara, I.M., Geiger, O. (2022) Specialized acyl carrier protein used by serine palmitoyltransferase to synthesize sphingolipids in *Rhodobacteria*. Front Microbiol 13: 961041. doi: 10.3389/fmicb.2022.961041

Pereira, T.C., de Barros, P.P., de Oliveira Fugisaki, L.R., Rossoni, R.D., de Camargo Ribeiro, F., de Menezes, R., et al. (2018) Recent advances in the use of *Galleria mellonella* model to study immune responses against human pathogens. J Fungi 4: 128

Price, M.N., Wetmore, K.M., Waters, R.J., Callaghan, M., Ray, J., Liu, H., et al. (2018) Mutant phenotypes for thousands of bacterial genes of unknown function. Nature 557: 503–509. doi: 10.1038/s41586-018-0124-0.

Sambrook, J., MacCullum, P., and Russell, D. (2001) Molecular Cloning: A Laboratory Manual, 3rd edn. Plainview, NY, USA: Cold Spring Harbor Laboratory Press.

Simon, R., Priefer, U., and Pühler, A. (1983) A broad host range mobilization system for in vivo genetic engineering: transposon mutagenesis in gram-negative bacteria. Biotechnology 1: 784– 791.

Sims, R.P.A., Larose, J.A.G. (1961) The use of iodine vapor as a general detecting agent in the thin layer chromatography of lipids. J Am Oil Chem Soc 39: 232.

Stankeviciute, G., Guan, Z., Goldfine, H., and Klein, E.A. (2019) *Caulobacter crescentus* adapts to phosphate starvation by synthesizing anionic glycoglycerolipids and a novel glycosphingolipid. mBio 10: e00107–19. Doi: 10.1128/mBio.00107-19.

Stankeviciute, G., Tang, P., Ashley, B., Chamberlain, J.D., Hansen, M.E.B., Coleman, A., D′Emilia, R., Fu, L., Mohan, E.C., Nguyen, H., Guan, Z., Campopiano, D.J., and Klein, E.A. (2022) Convergent evolution of bacterial ceramide synthesis. Nature Chem Biol 18:305–312. Doi: 10.1038/s41589-021-00948-7.

Thanbichler, M., Iniesta, A.A., and Shapiro, L. (2007) A comprehensive set for vanillate-and xylose-inducible gene expression in *Caulobacter crescentus*. Nucleic Acids Res 35:e137.

Zbierzak, A.M., Dörmann, P., and Hölzl G. (2011) Analysis of lipid content and quality in Arabidopsis plastids. In: Chloroplast Research in Arabidopsis: Methods and Protocols, Volume II, Methods in Molecular Biology, vol. 775. R.P. Jarvis (ed). Berlin: Springer, pp. 421–426. doi: 10.1007/978-1-61779-237-3_22

Zik, J.J., Yoon, S.H, Guan, Z., Stankeviciute Skidmore, G., Gudoor, R.R., Davies, K.M., Deutschbauer, A.M., Goodlett, D.R., Klein, E.A., Ryan, K.R. (2022). *Caulobacter* lipid A is conditionally dispensable in the absence of *fur* and in the presence of anionic sphingolipids. Cell Rep 39:110888. doi: 10.1016/j.celrep.2022.110888.

