## Supplemental Data for "Genes required for the formation of virulence-provoking bacterial sphingolipids"

### Supplementary Results

*Bioinformatic analyses of CC_1168 - CC_1152 candidate genes for sphingolipid biosynthesis and transport*

In order to identify candidate genes that might be involved in the conversion of ceramide to PSphLs, we analyzed the genomic region comprising CC_1168 - CC_1152 because many of those genes were linked to ceramide-producing genes through the cofitness results of their respective mutants.

CC_1168 displays the PFAM domain PF06835 which belongs to the LptC family. Following extraction of LPS from the IM, LptC was shown to participate in the transport of LPS to the OM of Gram-negative bacteria (Okuda *et al*., 2016). In *C. crescentus*, CC_1168 might be needed for PSphL transport from the IM to the OM. When predicting transmembrane protein topology with a hidden Markov model (TMHMM) (Krogh *et al*., 2001) or Phobius (Käll *et al*., 2007), CC_1168 has a possible transmembrane helix close to its N-terminus (Fig. S2). Fitness browser suggests that CC_1168 mutants display cofitness with mutants deficient in CC_1167, CC_1161, CC_1160, CC_1159, CC_1158, CC_1156, and CC_1152 (Table S1).

CC_1167 has the InterPro domain IPR007554, suggesting that it might encode a CDP-glycerol glycerophosphotransferase or glycosyltransferase. A ceramide-UTP-glucuronosyltransferase-deficient mutant of *Zymomonas mobilis* accumulates ceramide-phospho-glycerol (Okino *et al*., 2020). There are good homologs (ZMO1734; E = 4e-83 and Swit_4739; E = 8e-99) of CC_1167 encoded by the *Z. mobilis* and *Sphingomonas wittichii* genomes, respectively. But presently their exact function is unknown.

CC_1166 has the InterPro domain IPR002797, suggesting that it might encode an oligosaccharide transport membrane protein (flippase) involved in synthesis of the O-antigen of LPS. According to TMHMM and Phobius, CC_1166 has nine transmembrane helices (Fig. S2).

CC_1165 encodes the specialized acyl-ACP synthetase AasR which specifically acylates the special acyl carrier protein AcpR (Padilla-Gómez *et al*., 2022).

CC_1164 is predicted to encode an NADPH-dependent epimerase/dehydrogenase and is probably responsible for the reduction of the 3-oxo group in 3-oxo-sphinganine. However, recently it was suggested that CC_1164 would encode a dehydrogenase, also termed CerR (Stankeviciute *et al.*, 2022), that reduces *N*-acylated 3-oxo-sphinganine (3-oxo-dihydroceramide) and that *N*-acylation of 3-oxo-sphinganine by CC_1154, also termed bCerS (Stankeviciute *et al.*, 2022), would precede the reduction step (Stankeviciute *et al*., 2022).

CC_1163 is the specialized acyl carrier protein AcpR, required for efficient SphL biosynthesis in *Rhodobacteria* (Padilla-Gómez *et al*., 2022).

CC_1162 encodes an Spt that preferentially uses palmitoyl-AcpR as acyl donor during 3-oxo-sphinganine synthesis (Padilla-Gómez *et al*., 2022).

There are homologs of CC_1161 in α-, β-, γ-, and δ-proteobacteria and they contain the InterPro domain IPR015222 of "mitochondrial cytidylyltransferase phosphatides" (Tam41/Mmp7) (CDP-DAG synthase) which catalyzes CDP-DAG formation from phosphatidic acid. Fitness browser suggests that CC_1161-deficient mutants display cofitness with mutants deficient in CC_1168, CC_1160, CC_1159, CC_1158, CC_1156, CC_1152, CC_1385 (predicted fructose-1,6-bisphosphatase) (Table S1), and CC_0001 (putative pyruvate water, dikinase/phospenolpyruvate synthetase). It is remarkable that CC_1385 as well as CC_0001 seem to encode enzymes participating in gluconeogenesis. Based on mass spectrometric data, Zik *et al*. (2022) propose that CC_1161 attaches a glycerate residue to ceramide-1-phosphate.

CC_1160 was annotated as sphingosine/diacylglycerol (DAG) kinase. TMHMM predicts a transmembrane helix in CC_1160 with an N-terminal cytoplasmic and a C-terminal periplasmic domain (Fig. S2), while Phobius predicts two transmembrane helices (Fig. S2). Fitness browser suggests that CC_1160 mutants display cofitness with mutants deficient in CC_1168, CC_1167, CC_1161, CC_1159, CC_1158, and CC_1156 (Table S1). Recent experimental work indicates that CC_1160 is a novel ceramide kinase that can phosphorylate ceramide as well as DAG (Zik *et al*., 2022; Dhakephalkar *et al.*, 2023).

CC_1159 has the PFAM domain PF01066 which suggests displacement of CMP from a CDP-alcohol in order to condense with a second alcohol. Both TMHMM and Phobius suggests an N-terminal cytoplasmic domain and three transmembrane helices for CC_1159 (Fig. S2). Based on mass spectrometric data, Zik *et al*. (2022) propose that CC_1159 is a 2-phosphoglycerate transferase.

CC_1158 has a phosphoesterase InterPro domain IPF004843 similar to that of calcineurin type ApaH. These phosphoesterases are specific for 3′, 5′-cAMP. Fitness browser suggests cofitness of mutants affected in CC_1168, CC_1167, CC_1161, CC_1160, CC_1159, CC_1156, or CC_1152 with a mutant deficient in CC_1158 (Table S1).

CC_1157 has a histidine triad motif (HIT) consisting of HXHXHXX, where X is a hydrophobic amino acid. Proteins with HIT domains form a superfamily of nucleotide hydrolases and transferases which act on the α-phosphate of ribonucleotides.

CC_1156 contains a PFAM domain PF03739 (LptG/LptF) and codes for an integral membrane protein (TMHMM and Phobius predicts 6 transmembrane helices) (Fig. S2) which might participate in the export of LPS or PSphLs. Fitness browser suggest that CC_1156 mutants display cofitness with mutants deficient in CC_1168, CC_1166, CC_1161, CC_1160, CC_1159, CC_1158, and CC_1152 (Table S1).

CC_1155 also contains the PFAM domain PF03739 (LptG/LptF) and codes for an integral membrane protein (TMHMM and Phobius predicts six transmembrane helices) (Fig. S2) which also might participate in the export of LPS or PSphLs.

CC_1154 encodes the *N*-acyltransferase bCerS that can convert 3-oxo-sphinganine or sphinganine to their respective *N*-acylated products (Stankeviciute *et al*., 2022).

CC_1153 is similar to the InterPro domain IPR029004 of GTP:adenosylcobinamide-phosphate guanilyltransferase (CobY) and to the biosynthesis protein for molybdopterine-guanine A (MobA) IPR025877. KEGG suggests that homologs are annotated as 2-phospho-L-lactate guanilyltransferase. This family is represented by CofC (Bashiri *et al*., 2019), a nucleotidyltransferase that participates in the biosynthesis of coenzyme F420.

CC_1152 has PFAM domains PF00483, PF1128, and PF2804 of nucleotidyltransferases. Many enzymes that transfer nucleotides to phosphosugars are included in this family. Fitness browser suggests that CC_1152-deficient mutant shows cofitness with mutants deficient in CC_1168, CC_1166, CC_1161, CC_1159, CC_1158, CC_1385 (predicted fructose-1,6-bisphosphatase), and CC_0001 (putative pyruvate, water dikinase/phosphoenolpyruvate synthetase) (Table S1). Therefore, CC_1385 and CC_0001 seem to encode two critical gluconeogenesis enzymes.

In summary, homologs that encode transport proteins (CC_1168, CC_1166, CC_1156, CC_1155) might contribute to forming a complex transport system that moves PSphLs from their site of synthesis in the IM to their final destination in the outer layer of the OM in an analogous way to that in which LPS is transported. However, we did not investigate PSphL transport in more detail in this work. Instead, we focused on studying genes potentially coding for biosynthetic enzymes (CC_1161, CC_1160, CC_1159, CC_1158, CC_1153, CC_1152) that participate in PSphL biosynthesis.

### References

Bashiri, G., Antoney, J., Jirgis, E.N.M., Shah, M.V., Ney, B., Copp, J. et al. (2019) A revised biosynthetic pathway for the cofactor F420 in prokaryotes. *Nat Commun* **10:** 1558. doi: 10.1038/s41467-019-09534-x

Dhakephalkar, T., Stukey, G.J., Guan, Z., Carman, G.M., and Klein, E.R. (2023) Characterization of an evolutionary distinct bacterial ceramide kinase from *Caulobacter crescentus*. *J Biol Chem* **299:** 104894. doi: 10.1016/j.jbc.2023.104894

Käll, L., Krogh, A., & Sonnhammer, E. L. (2007) Advantages of combined transmembrane topology and signal peptide prediction--the Phobius web server. *Nucleic Acids Res* **35**(Web Server issue): W429–W432. DOI: 10.1093/nar/gkm256

Krogh, A., Larsson, B., von Heijne, G., & Sonnhammer, E. L. (2001) Predicting transmembrane protein topology with a hidden Markov model: application to complete genomes. *J Mol Biol* **305**(3): 567–580. doi: 10.1006/jmbi.2000.4315.PMID: 11152613

Okino, N., Li, M., Qu, Q., Nakagawa, T., Hayashi, Y., Matsumoto, M., Ishibashi, Y., and Ito, M. (2020). Two bacterial glycosphingolipid synthases responsible for the synthesis of glucuronosylceramide and α-galactosylceramide. *J Biol Chem* **295:** 10709–10725.

Okuda, S., Sherman, D. J., Silhavy, T. J., Ruiz, N., and Kahne, D. (2016) Lipopolysaccharide transport and assembly at the outer membrane: the PEZ model. *Nature Rev Microbiol* **14:** 337–345.

Padilla-Gómez, J., Olea-Ozuna, R.J., Contreras-Martínez, S., Morales-Tarré, O., García-Soriano, D.A. Sahonero-Canavesi, D.X., Poggio, S., Encarnación-Guevara, S., López-Lara, I.M., Geiger, O. (2022) Specialized acyl carrier protein used by serine palmitoyltransferase to synthesize sphingolipids in *Rhodobacteria*. *Front Microbiol* **13:** 961041. doi: 10.3389/fmicb.2022.961041

Stankeviciute, G., Tang, P., Ashley, B., Chamberlain, J.D., Hansen, M.E.B., Coleman, A., D´Emilia, R., Fu, L., Mohan, E.C., Nguyen, H., Guan, Z., Campopiano, D.J., and Klein, E.A. (2022) Convergent evolution of bacterial ceramide synthesis. *Nature Chem Biol* **18:**305-312. Doi: 10.1038/s41589-021-00948-7.

Zik, J.J., Yoon, S.H, Guan, Z., Stankeviciute Skidmore, G., Gudoor, R.R., Davies, K.M., Deutschbauer, A.M., Goodlett, D.R., Klein, E.A., Ryan, K.R. (2022). *Caulobacter* lipid A is conditionally dispensable in the absence of *fur* and in the presence of anionic sphingolipids. *Cell Rep* **39:**110888. doi: 10.1016/j.celrep.2022.110888.

### Supplementary Figures

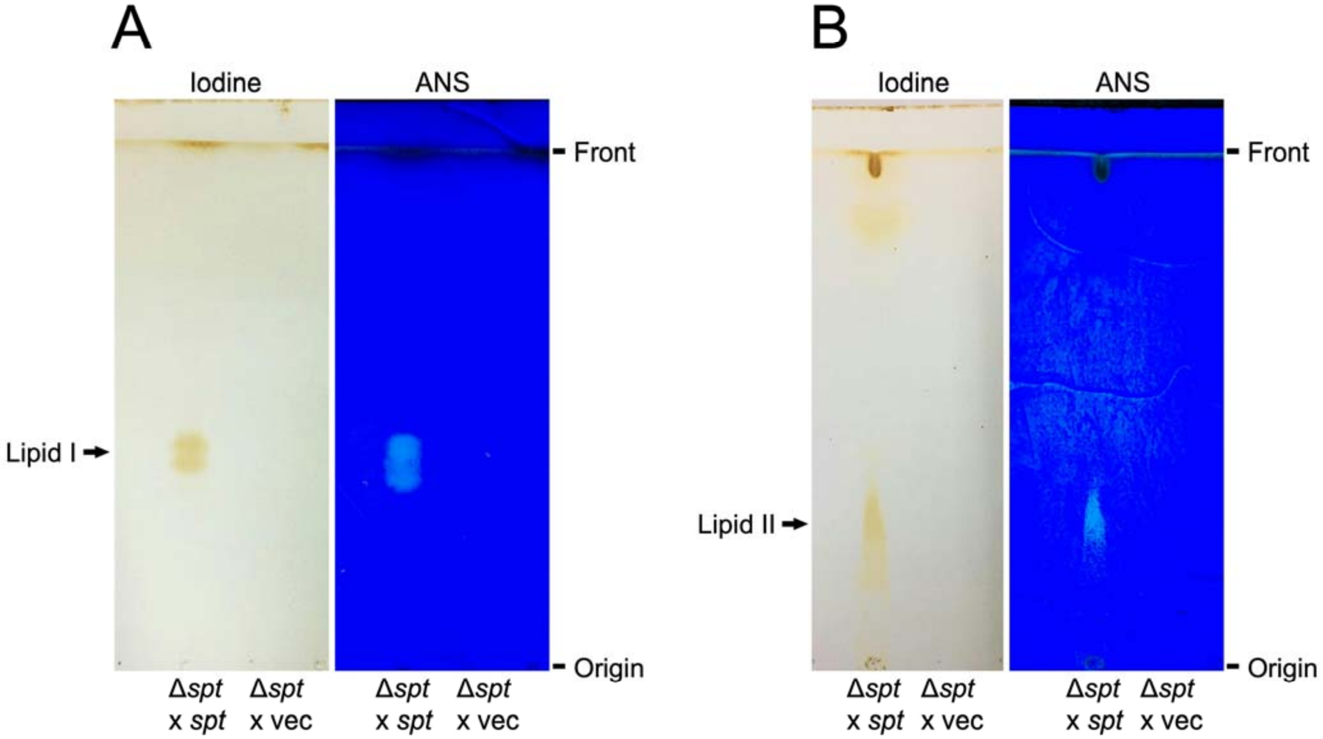


**Fig. S1.** Fractions in Lipid I and Lipid II after purification by TLC. Lipid extracts from large cultures (2 l) of the *spt*-deficient mutant harboring the *spt* gene *in trans* (Δ*spt* x *spt*) or of the *spt*-deficient mutant harboring the empty vector pRXMCS-2 (Δ*spt* x vec) of *C. crescentus* were obtained, separated by preparative TLC, and visualized on exposure to iodine vapor. From areas that contained Lipid I or II, silica gel was scraped, lipids were extracted, and aliquots were reanalyzed by TLC in order to assess their purity. As visualized after iodine or ANS staining, fractions enriched in Lipid I (panel A) or Lipid II (panel B) were obtained from the SphL-producing Δ*spt* x *spt* strain while no analogous lipids were observed following the same extraction procedures carried out on the Δ*spt* x vec strain of *C. crescentus.*

CC_1168

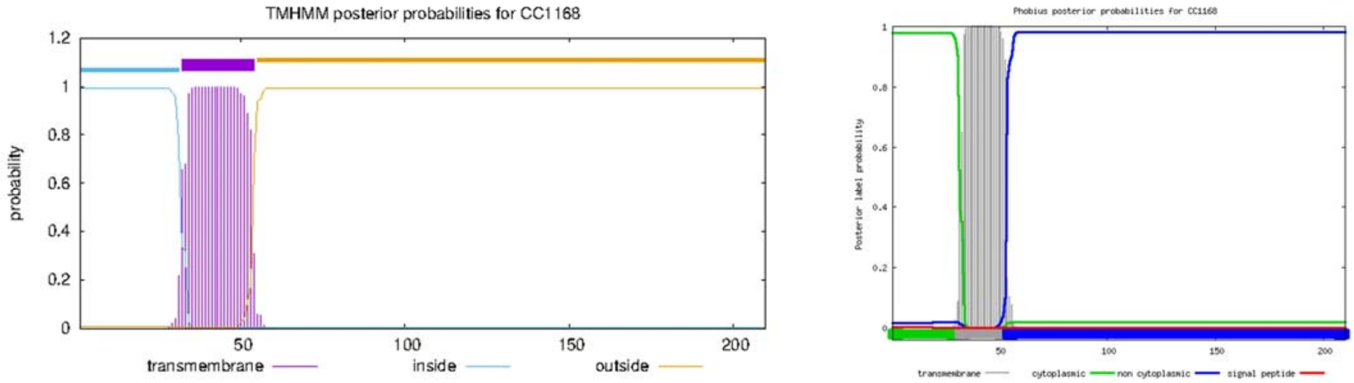


CC_1166

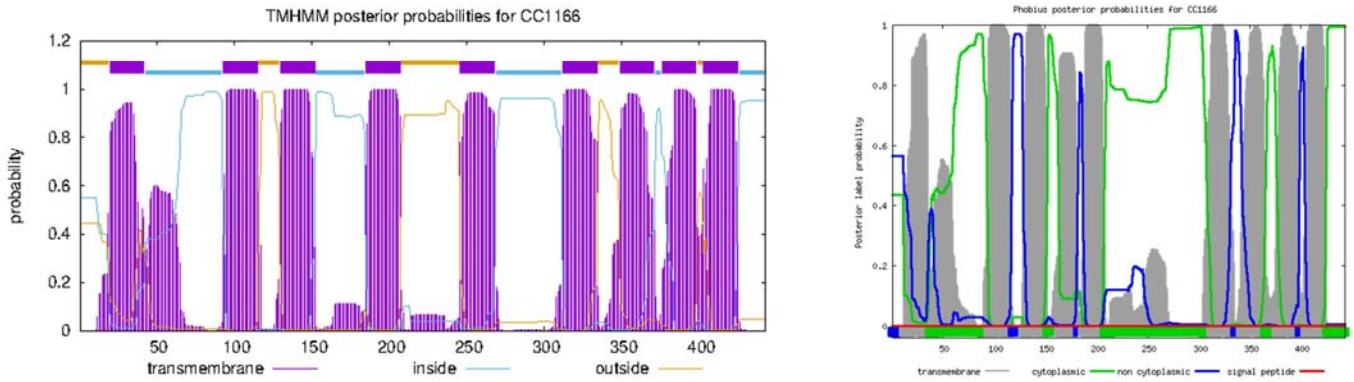


CC_1160

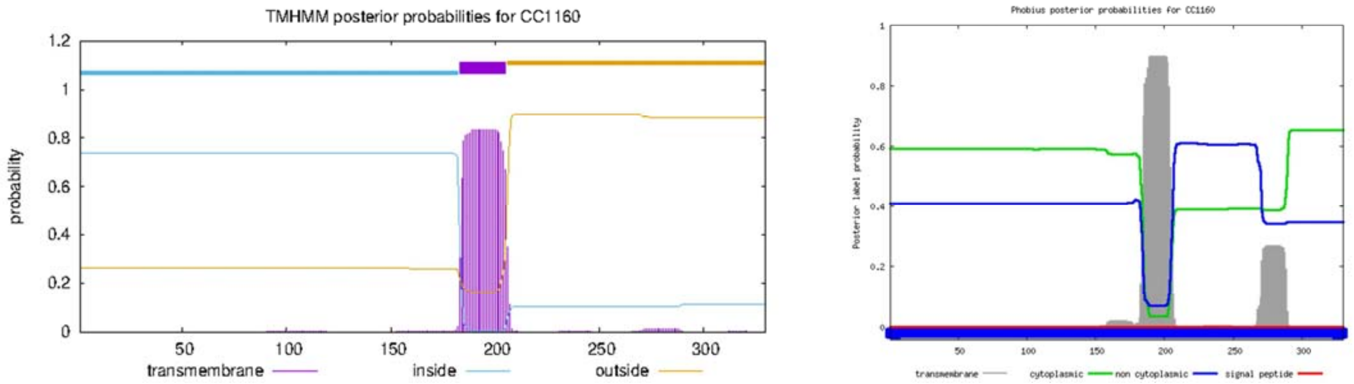

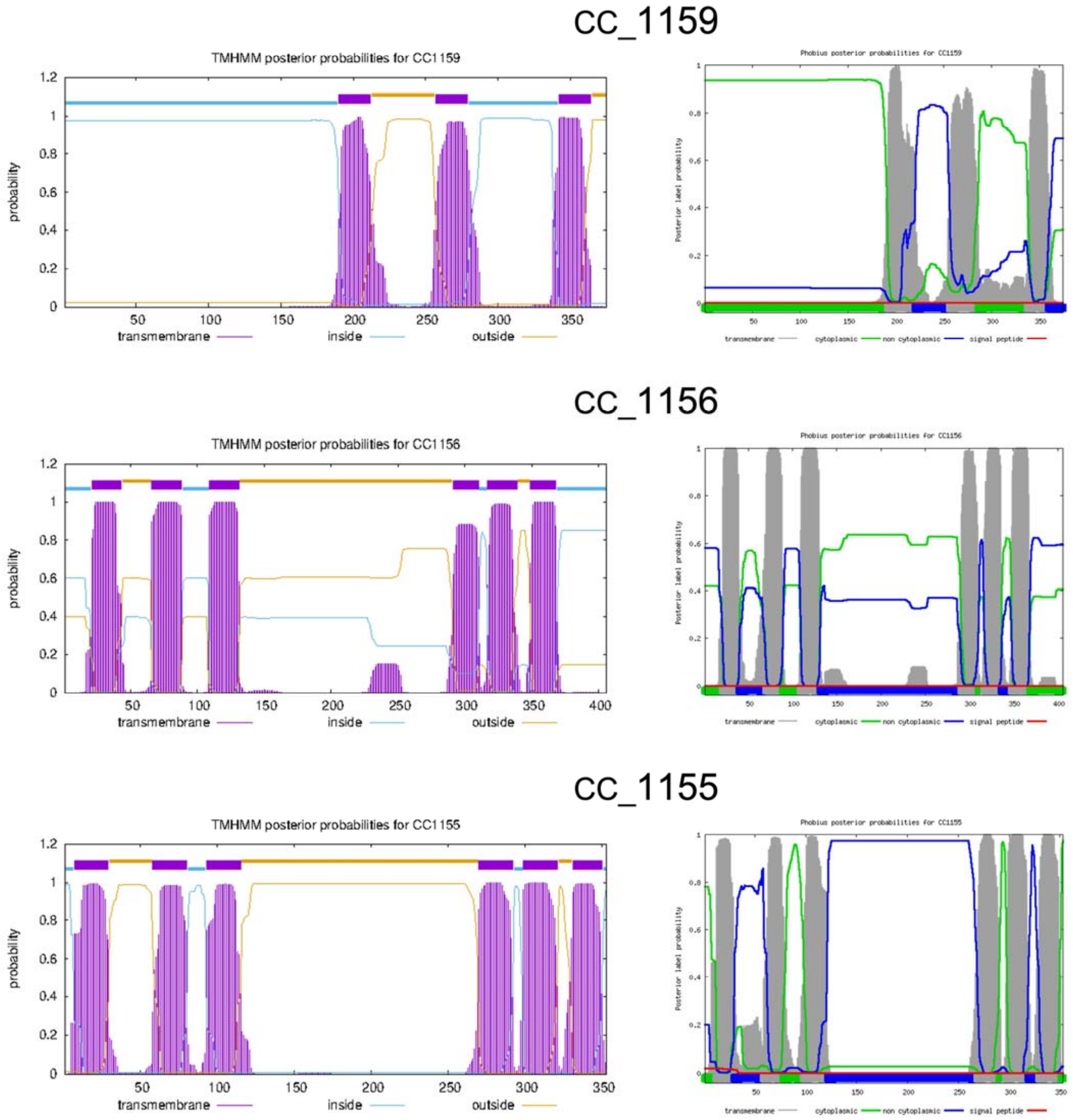


**Fig. S2.** Prediction of transmembrane helices for CC_1168, CC_1166, CC_1160, CC_1159, CC_1156, and CC_1155 using the bioinformatic programs TMHMM 2.0 (left) and Phobius (right).

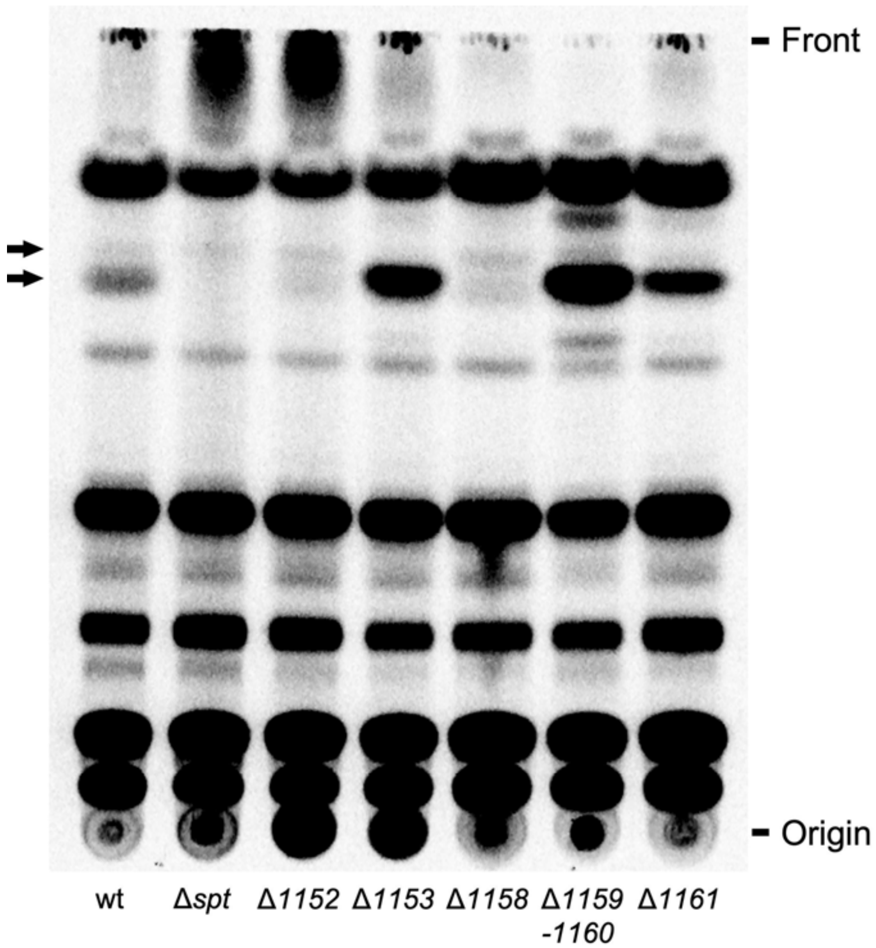


**Fig. S3.** Dihydroceramide profiles of mutants deficient in genes required for PSphL formation. Different *C. crescentus* strains [wild-type strain (wt), *spt*-deficient mutant (Δ*spt*), mutant SPG14 deficient in CC_1152 (Δ*1152*), mutant SPG15 deficient in CC_1153 (Δ*1153*), mutant SPG11 deficient in CC_1158 (Δ*1158*), mutant SPG09 deficient in CC_1159-1160 (Δ*1159-1160*), and mutant SPG18 deficient in CC_1161 (Δ*1161*)] were cultured in complex medium in the presence of $^{14}$C-acetate. After harvesting cells, lipids were extracted, separated by TLC in chloroform/methanol/ammonium hydroxide (40:10:1) and developed chromatograms were analyzed by phosphorimaging. Arrows indicate dihydroceramides formed by *C. crescentus*.

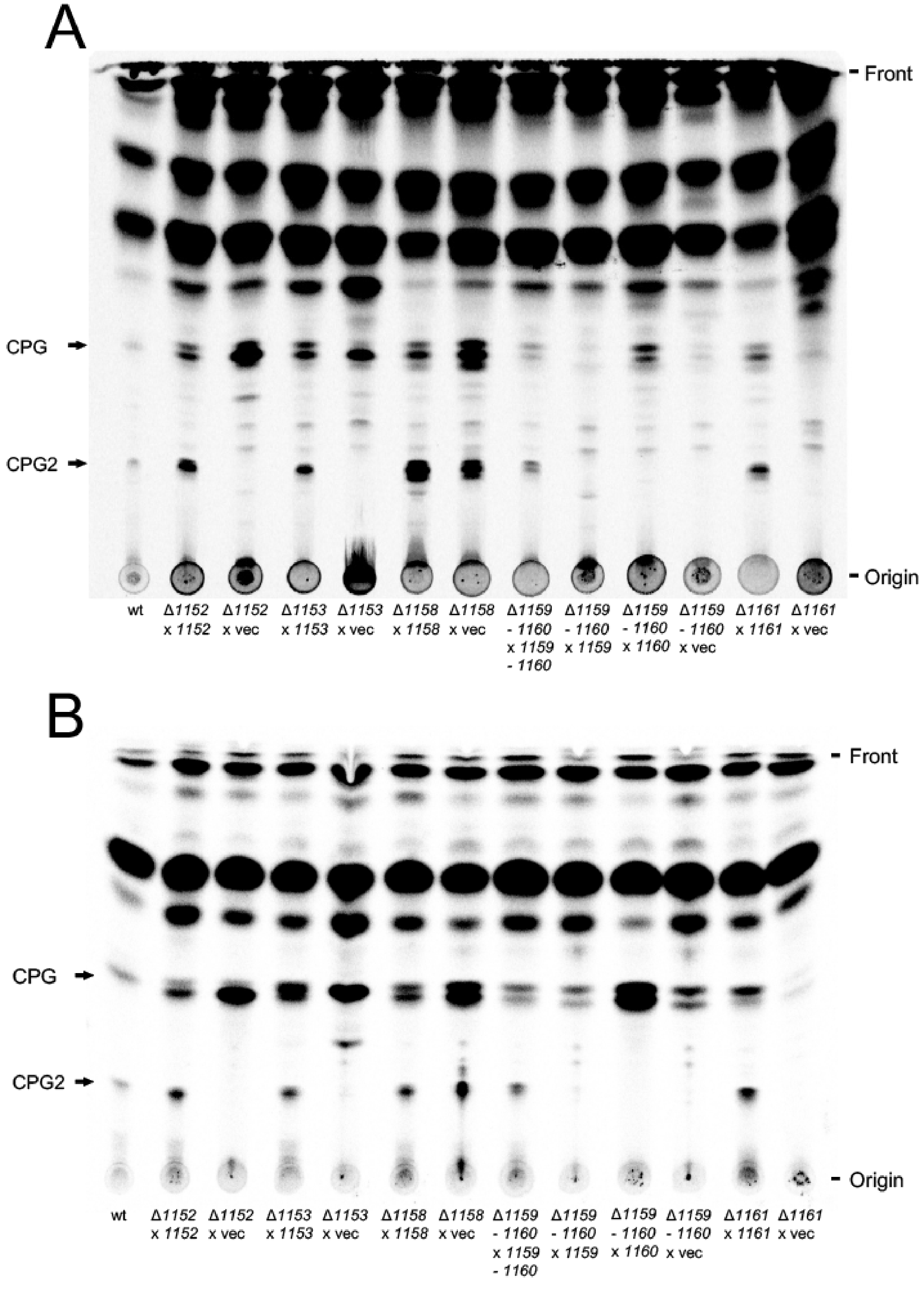
A
Front
CPG
CPG2
Origin
wt
Δ1152 x 1152
Δ1152 x vec
Δ1153 x 1153
Δ1153 x vec
Δ1158 x 1158
Δ1158 x vec
Δ1159 - 1160 x 1159 - 1160
Δ1159 - 1160 x 1159
Δ1159 - 1160 x 1160
Δ1159 - 1160 x vec
Δ1161 x 1161
Δ1161 x vec
B
Front
CPG
CPG2
Origin
wt
Δ1152 x 1152
Δ1152 x vec
Δ1153 x 1153
Δ1153 x vec
Δ1158 x 1158
Δ1158 x vec
Δ1159 - 1160 x 1159 - 1160
Δ1159 - 1160 x 1159
Δ1159 - 1160 x 1160
Δ1159 - 1160 x vec
Δ1161 x 1161
Δ1161 x vec

**Fig. S4.** Complementation of *C. crescentus* mutants altered in the formation of PSphL (CPG and CPG2) by their native genes *in trans*. Mutants of *C. crescentus* deficient in CC_1152, CC_1153, CC_1158, CC_1159-1160, or CC_1161 carrying the respective intact gene in the xylose-inducible plasmid pBXMCS-2 (Δ*1152* x *1152*, Δ*1153* x *1153*, Δ*1158* x *1158*, Δ*1159-1160* x *1159*, Δ*1159-1160* x *1160*, Δ*1159-1160* x *1159-1160*, Δ*1161* x *1161*) or carrying the empty pBXMCS-2 vector (vec) *in trans* were radiolabeled with $^{14}$C-acetate (A) or $^{33}$P-phosphate (B) for 16 h. At the end of the labeling period, cells were harvested, lipids were extracted, separated by TLC in chloroform/methanol/acetic acid/water (8:3:2:1) and developed chromatograms were subjected to autoradiography. Arrows indicate PSphLs (CPG and CPG2) formed by *C. crescentus*.

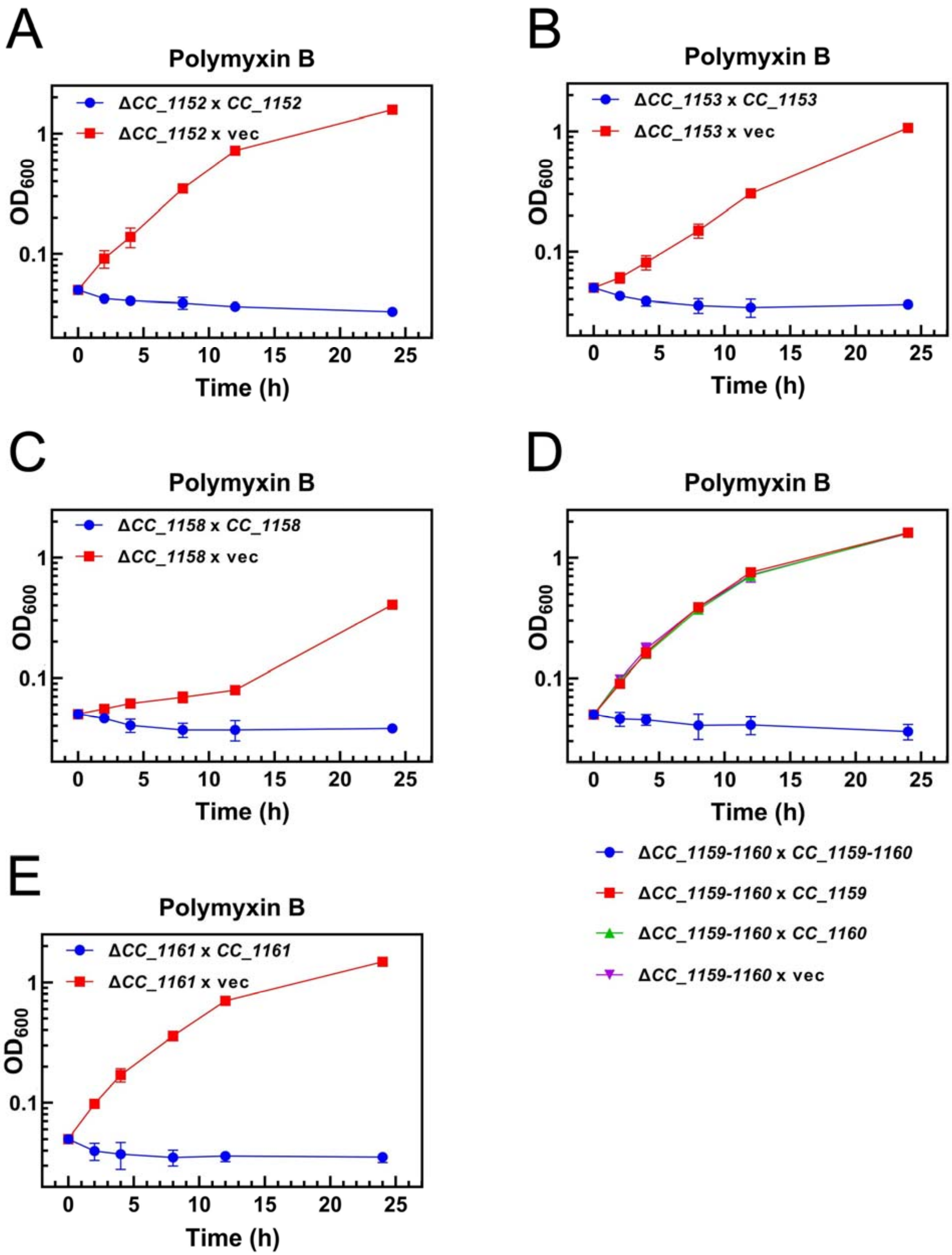
A
Polymyxin B
ΔCC_1152 x CC_1152
ΔCC_1152 x vec
OD600
Time (h)
B
Polymyxin B
ΔCC_1153 x CC_1153
ΔCC_1153 x vec
OD600
Time (h)
C
Polymyxin B
ΔCC_1158 x CC_1158
ΔCC_1158 x vec
OD600
Time (h)
D
Polymyxin B
OD600
Time (h)
ΔCC_1159-1160 x CC_1159-1160
ΔCC_1159-1160 x CC_1159
ΔCC_1159-1160 x CC_1160
ΔCC_1159-1160 x vec
E
Polymyxin B
ΔCC_1161 x CC_1161
ΔCC_1161 x vec
OD600
Time (h)

**Fig. S5.** Complementation of *C. crescentus* mutants altered in the PSphL formation by their native genes *in trans* restores sensitivity to polymyxin B. Growth ($OD_{600}$) of *C. crescentus* mutants deficient in CC_1152 harboring the CC_1152-expressing plasmid (Δ*1152* x *1152*) or the empty vector pBXMCS-2 (Δ*1152* x vec) (A), mutants deficient in CC_1153 harboring the CC_1153-expressing plasmid (Δ*1153* x *1153*) or the empty vector pBXMCS-2 (Δ*1153* x vec) (B), mutants deficient in CC_1158 harboring the CC_1158-expressing plasmid (Δ*1158* x *1158*) or the empty vector pBXMCS-2 (Δ*1158* x vec) (C), mutants deficient in CC_1159-1160 harboring the CC_1159-1160-expressing plasmid (Δ*1159-1160* x *1159-1160*), the CC_1159-expressing plasmid (Δ*1159-1160* x *1159*), the CC_1160-expressing plasmid (Δ*1159-1160* x *1160*), or the empty vector pBXMCS-2 (Δ*1159-1160* x vec) (D), and mutants deficient in CC_1161 harboring the CC_1161-expressing plasmid (Δ*1161* x *1161*) or the empty vector pBXMCS-2 (Δ*1161* x vec) (E). Data and bars represent the average and standard errors obtained from at least three independent experiments. Note that growth curves for strains Δ*1159-1160* x *1159*, Δ*1159-1160* x *1160*, and Δ*1159-1160* x vec overlap in (D).

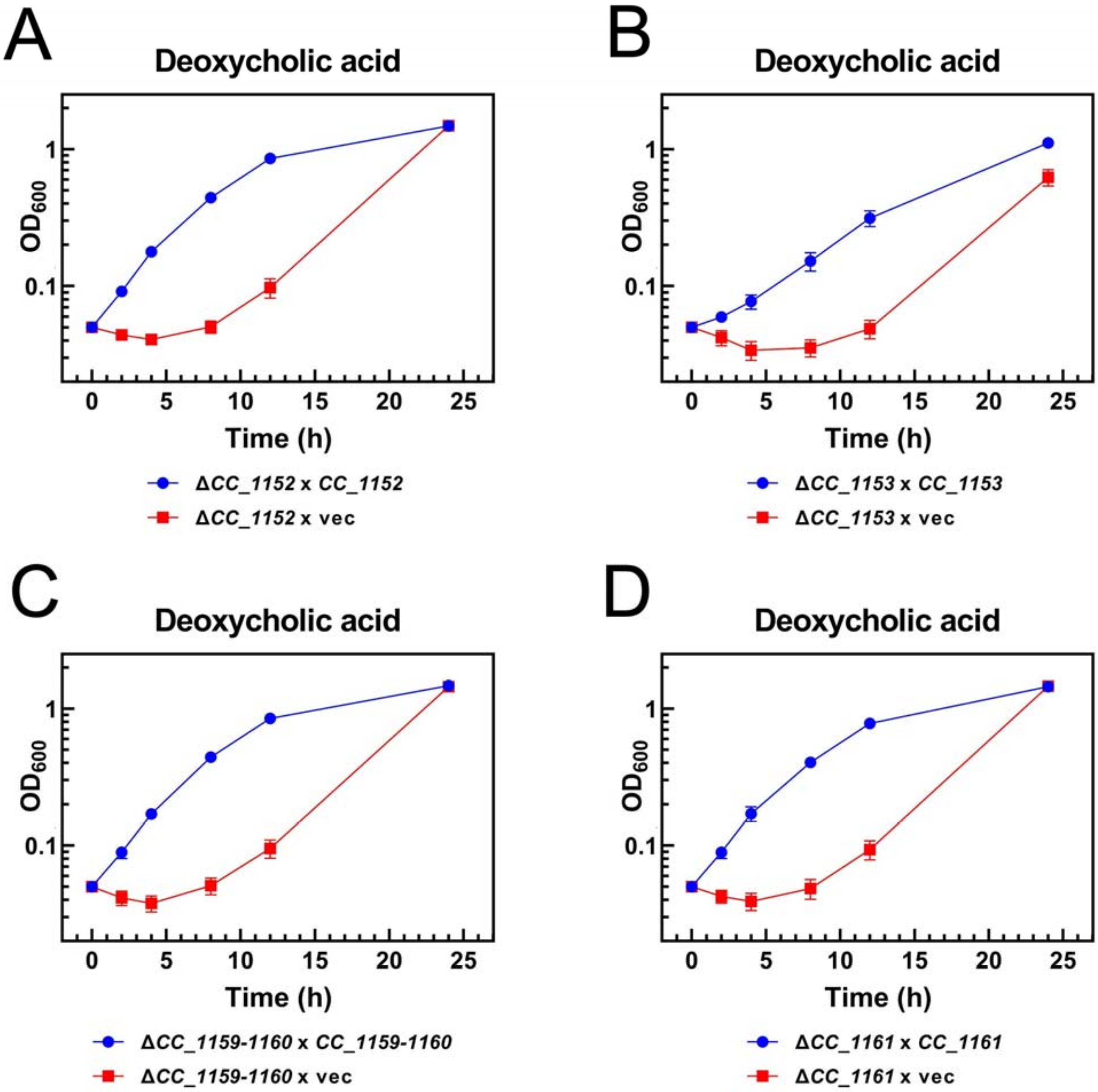


**Fig. S6.** Genes for PSphL formation are required for resistance of *C. crescentus* to deoxycholate. Growth ($OD_{600}$) of different strains of *C. crescentus* [mutant SPG14 deficient in CC_1152 expressing intact CC_1152 *in trans* (Δ*1152 x 1152*) or mutant harboring the empty vector (Δ*1152 x* vec)(A), mutant SPG15 deficient in CC_1153 expressing intact CC_1153 *in trans* (Δ*1153 x 1153*) or mutant harboring the empty vector (Δ*1153 x* vec)(B),, mutant SPG09 deficient in CC_1159/CC_1160 expressing intact CC_1159/CC_1160 *in trans* (Δ*1159-1160 x 1159/1160*) or mutant harboring the empty vector (Δ*1159-1160 x* vec)(C), and mutant SPG18 deficient in CC_1161 expressing intact CC_1161 *in trans* (Δ*1161 x 1161*) or mutant harboring the empty vector (Δ*1161 x* vec)(D)] was determined at 30ºC on complex medium in the presence of deoxycholate (1 mg/ml). Data and bars represent the average and standard errors obtained from at least three independent experiments.

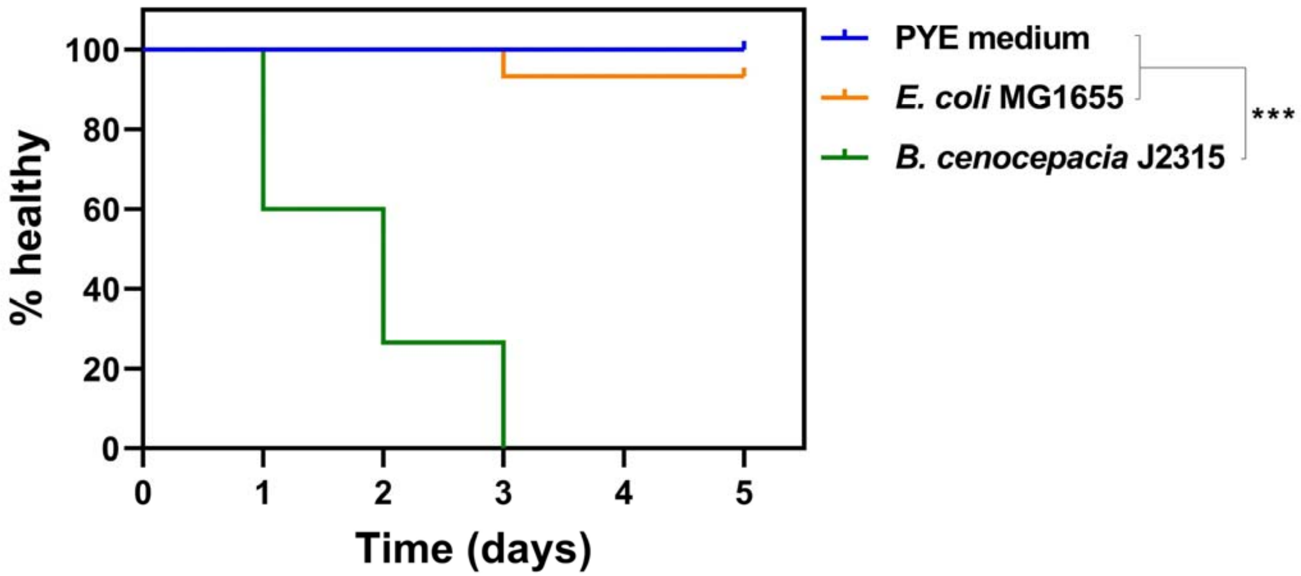


**Fig. S7.** Healthspan assays of *G. mellonella* larvae was determined with Kaplan-Meier survival analysis up to 5 days after inoculation with PYE medium, *E. coli* MG1655 ($10^5$ CFU), or *B. cenocepacia* J2315 ($10^5$ CFU). Survival curves are a representative cohort (n = 15) of at least three biological replicates (Mantel-Cox test for statistics, ***P<0.001).

**Table S1.** Cofitness of mutants affected in genes putatively involved in PSphL biosynthesis and transport in *C. crescentus*. Cofitness data were taken from Price *et al*. (2018). Mutants in CC_1157 do not show high cofitness values with other mutants of the cluster. For CC_1155 and CC_1153 no data have been reported. CC_1165, CC_1164, CC_1163, CC_1162, or CC_1154 are indicated as *aasR*, *cerR*, *acpR*, *spt*, or *cerS*, respectively.

| ***ΔCC_*** | ***1168*** | ***1167*** | ***1166*** | ***aasR*** | ***cerR*** | ***acpR*** | ***spt*** | ***1161*** | ***1160*** | ***1159*** | ***1158*** | ***1157*** | ***1156*** | ***1155*** | ***cerS*** | ***1153*** | ***1152*** | ***1385*** |
|---|---|---|---|---|---|---|---|---|---|---|---|---|---|---|---|---|---|---|
| ***1168*** | | 0.47 | | 0.59 | 0.62 | | 0.52 | 0.66 | 0.64 | 0.78 | 0.60 | | 0.67 | | 0.62 | | 0.63 | |
| ***1167*** | 0.47 | | | 0.64 | 0.51 | 0.57 | 0.57 | | 0.49 | 0.54 | 0.66 | | | | 0.64 | | | |
| ***1166*** | | | | | | | | | | | | | 0.62 | | | | 0.61 | 0.51 |
| ***aasR*** | 0.59 | 0.64 | | | 0.86 | 0.82 | 0.92 | | 0.79 | 0.71 | 0.68 | | 0.39 | | 0.94 | | | |
| ***1164*** | 0.62 | 0.51 | | 0.86 | | 0.69 | 0.80 | | 0.86 | 0.72 | 0.65 | | 0.49 | | 0.86 | | | |
| ***acpR*** | | 0.57 | | 0.82 | 0.69 | | 0.88 | | 0.60 | 0.52 | 0.60 | | | | 0.73 | | | |
| ***spt*** | 0.52 | 0.57 | | 0.92 | 0.80 | 0.88 | | | 0.67 | 0.60 | 0.60 | | | | 0.83 | | | |
| ***1161*** | 0.66 | | | | | | | | 0.44 | 0.79 | 0.48 | | 0.61 | | 0.42 | | 0.82 | 0.67 |
| ***1160*** | 0.64 | 0.49 | | 0.79 | 0.86 | 0.60 | 0.67 | 0.44 | | 0.75 | 0.62 | | 0.49 | | 0.80 | | | 0.53 |
| ***1159*** | 0.78 | 0.54 | | 0.71 | 0.72 | 0.52 | 0.60 | 0.79 | 0.75 | | 0.72 | | 0.60 | | 0.75 | | 0.74 | |
| ***1158*** | 0.60 | 0.66 | | 0.68 | 0.65 | 0.60 | 0.60 | 0.48 | 0.62 | 0.72 | | | 0.48 | | 0.73 | | 0.46 | |
| ***1157*** | | | | | | | | | | | | | | | | | | |
| ***1156*** | 0.67 | | 0.62 | 0.39 | 0.49 | | | 0.61 | 0.49 | 0.60 | 0.48 | | | | 0.43 | | 0.63 | |
| ***1155*** | | | | | | | | | | | | | | | | | | |
| ***1154*** | 0.62 | 0.64 | | 0.94 | 0.86 | 0.73 | 0.83 | 0.42 | 0.80 | 0.75 | 0.73 | | 0.43 | | | | | |
| ***1153*** | | | | | | | | | | | | | | | | | | |
| ***1152*** | 0.63 | | 0.61 | | | | | 0.82 | | 0.74 | 0.46 | | 0.63 | | | | | 0.71 |
| ***1385*** | | | 0.51 | | | | | 0.67 | | 0.53 | | | | | | | 0.71 | |

**Table S2**. Construction of *C. crescentus* knock-out mutants in potential sphingolipid biosynthesis genes.

| *C. crescentus* mutant | Mutant characteristics and oligonucleotide primers used for construction | Restriction site | PCR product (bp) |
|---|---|---|---|
| **SPG14** | ***cc_1152* in frame deletion** | | |
| | primer pair for upstream region: | | |
| | CAAAAAGCTTCCAAGGCCGCCGCCGACATCG | HindIII | |
| | CAAAGGATCCAACCGGCTGCATGACGTTCTGG | BamHI | 614 |
| | primer pair for downstream region: | | |
| | CAAAGGATCCCAGGAGGCCGAGGCGGTCTAGG | BamHI | |
| | CAAAGAATTCCCTGGCTGACACCCCTGACTGG | EcoRI | 618 |

In mutant SPG14 (Δ*1152*) a deletion of 729 bp, coding for amino acid residues 5-247 of a predicted nucleotidyltransferase family protein (253 amino acid residues in total), is replaced by a hexanucleotide providing a BamHI restriction site and coding for G and S.

| | | | |
|---|---|---|---|
| **SPG15** | ***cc_1153* in frame deletion** | | |
| | primer pair for upstream region: | | |
| | CAAAAAGCTTCCCCGCCGATGGTCTTGATGG | HindIII | |
| | CAAGGATCCACCCGCGAGGATCAGGGCCTTG | BamHI | 739 |
| | primer pair for downstream region: | | |
| | CAAAGGATCCCTGGTGGGCGAGGCGAAAAGC | BamHI | |
| | CAAAGAATTCAGGTCGCGCGCCTGTCGCTGC | EcoRI | 748 |

In mutant SPG15 (Δ*1153*) a deletion of 705 bp, coding for amino acid residues 44-278 of a predicted MobA-like NTP transferase domain protein (291 amino acid residues in total), is replaced by a hexanucleotide providing a BamHI restriction site and coding for G and S.

**SPG18** ***cc_1161*** **deletion**

primer pair for upstream region:

| | | |
|---|---|---|
| CAAAAAGCTTCCTTGAAGGCGGCGCTGGACG | HindIII | |
| CAAAGGATCCTGGGAGAGCTCGGCCGCGACG | BamHI | 843 |

primer pair for downstream region:

| | | |
|---|---|---|
| CAAAGGATCCCTGGAGAGAGCGGCGGCAAGC | BamHI | |
| CAAAGAATTCGAAGACTTGGCCAAGCGCCTGC | EcoRI | 834 |

In mutant SPG18 (Δ*1161*) a deletion of 813 bp, coding for amino acid residues 14-284 of a predicted cytosolic protein (882 amino acid residues in total), is replaced by a hexanucleotide providing a BamHI restriction site and coding for G and S.

**Table S3.** Oligonucleotides used for amplification of different sphingolipid biosynthesis genes. Sites for recognition by restriction enzymes are underlined.

| **Primer Sequences** | **(5´-3´)** |
|---|---|
| **Primers for expression plasmids** | |
| oLOP444 | AGGAATACATATGCAGCCGGTTAAGACCCTTATTCTC |
| oLOP445 | ACTGGGATCCCTAGACCGCCTCGGCCTCCTGG |
| oLOP446 | AGGAATACATATGGGTTCTGGAGGCCAACAAGGC |
| oLOP447 | ACTGGAATTCTTAGCCTTTGAAATGTAAAGGGCTTTTCGC |
| oLOP448 | AGGAATACATATGTCCATTTATCGCATCGCCCACC |
| oLOP449 | ACTGGGATCCTCAGGCGGCCTCGGGG |
| oLOP450 | AGGAATACATATGAGTAGTGAAGTTCAAAAAGGGCCG |
| oLOP451 | ACTGGGATCCTCATTTCGCCAGCCAGGACTG |
| oLOP452 | AGGAATACATATGCTTCGTCGTGCACGCCATC |
| oLOP453 | ACTGGGTACCTCATCCGACCAGGAACCGCAAG |
| oLOP454 | AGGAATACATATGAGCCGCCTGCGCGG |
| oLOP455 | ACTGGGTACCCTATGCGGCTTGCCGCCGC |

**Table S4.** Construction of different expression plasmids.

For details see Experimental Procedures in main text.

| Plasmid | ORFs amplified | Oligonucleotides used | Restricted with | Cloned into plasmid restricted with ( ) |
|---|---|---|---|---|
| pRJ14 | *cc_1152* | oLOP444/oLOP445 | NdeI/BamHI | pET17b (NdeI/BamHI) |
| pRJ15 | *cc_1153* | oLOP446/oLOP447 | NdeI/EcoRI | pET17b (NdeI/EcoRI) |
| pRJ16 | *cc_1158* | oLOP448/oLOP449 | NdeI/BamHI | pET17b (NdeI/BamHI) |
| pRJ17 | *cc_1159* | oLOP450/oLOP451 | NdeI/BamHI | pET17b (NdeI/BamHI) |
| pRJ18 | *cc_1160* | oLOP452/oLOP453 | NdeI/KpnI | pET17b (NdeI/KpnI) |
| pRJ19 | *cc_1159/cc_1160* | oLOP450/oLOP453 | NdeI/KpnI | pET17b (NdeI/KpnI) |
| pRJ20 | *cc_1161* | oLOP454/oLOP455 | NdeI/KpnI | pET17b (NdeI/KpnI) |
| pRJ21 | *cc_1152* | Recloned from pRJ14 | NdeI/EcoRI | pBXMCS-2 (NdeI/EcoRI) |
| pRJ22 | *cc_1153* | Recloned from pRJ15 | NdeI/EcoRI | pBXMCS-2 (NdeI/EcoRI) |
| pRJ23 | *cc_1158* | Recloned from pRJ16 | NdeI/EcoRI | pBXMCS-2 (NdeI/EcoRI) |
| pRJ24 | *cc_1159* | Recloned from pRJ17 | NdeI/EcoRI | pBXMCS-2 (NdeI/EcoRI) |
| pRJ25 | *cc_1160* | Recloned from pRJ18 | NdeI/EcoRI | pBXMCS-2 (NdeI/EcoRI) |
| pRJ26 | *cc_1159/cc_1160* | Recloned from pRJ19 | NdeI/EcoRI | pBXMCS-2 (NdeI/EcoRI) |
| pRJ27 | *cc_1161* | Recloned from pRJ20 | NdeI/EcoRI | pBXMCS-2 (NdeI/EcoRI) |
